# GREEN-DB: A framework for the annotation and prioritization of non-coding regulatory variants from whole-genome sequencing data

**DOI:** 10.1101/2020.09.17.301960

**Authors:** E Giacopuzzi, N Popitsch, JC Taylor

## Abstract

Non-coding variants have emerged as important contributors to the pathogenesis of human diseases, not only as common susceptibility alleles but also as rare high-impact variants. Despite recent advances in the study of regulatory elements and the availability of specialized data collections, the systematic annotation of non-coding variants from genome sequencing remains challenging. Here, we propose a new framework for the prioritization of non-coding regulatory variants that integrates information about regulatory regions with prediction scores and HPO-based prioritization. Firstly, we created a comprehensive collection of annotations for regulatory regions including a curated database of 2.4 million regulatory elements (GREEN-DB) annotated with controlled gene(s), tissue(s) and associated phenotype(s) where available. Secondly, we calculated a variation constraint metric and showed that constrained regulatory regions associate with disease-associated genes and essential genes from mouse knock-out screens. Thirdly, we compared 19 non-coding impact prediction scores providing suggestions for variant prioritization. Finally, we developed a VCF annotation tool (GREEN-VARAN) that can integrate all these elements to annotate variants for their potential regulatory impact. In our evaluation, we show that GREEN-DB can capture previously published disease-associated non-coding variants as well as identify additional candidate disease genes in WGS trio analyses.

## Introduction

The precise spatiotemporal control of gene expression plays a fundamental role in developmental processes and cellular functions and consequently, is essential in determining human phenotypes (1–3). Gene expression is controlled by the interaction of distal regulatory elements, such as enhancers and silencers, with proximal gene promoters, and is mediated by complex networks of transcription factors (TF) binding to these genomic regions (4–7). Sequence variants within these regulatory regions can alter TF binding and/or enhancer-promoter interactions, resulting in gene expression dysregulation and eventually disease (8–13). The contribution of regulatory regions in human diseases is also supported by a myriad of genome-wide association studies (GWAS), showing that most disease-risk variants lie in non-coding regions (14–16). In recent years, our knowledge about regulatory mechanisms and regulatory elements across the human genome has substantially improved due to a large number of genomic, epigenomic, and transcriptomics studies. Main functional elements in the human genome, such as enhancers, promoters, and TF binding sites, have been extensively mapped by large international collaborations like ENCODE (17, 18) and FANTOM5 (19, 20). Several dedicated resources have subsequently been developed, integrating and extending these datasets to generate a more detailed picture of regulatory elements (21–26). Meanwhile, the application of novel computational (26–29) and high-throughput screening methods (30–33) has substantially improved our understanding of how regulatory elements control their respective target genes while several *in-silico* methods have been developed to better predict the impact of non-coding regulatory variants (34–40). The increasing adoption of whole-genome (WGS) over whole-exome (WES) sequencing in disease studies now allows for the comprehensive investigation of human variants (41), including those affecting these regulatory regions. The accurate identification, interpretation, and prioritization of such WGS-derived variants requires standardized resources for their annotation in routine bioinformatics pipelines in order to identify likely pathogenic regulatory variants. Whilst there is a large variety of annotation methods and databases available for coding variants (42, 43), resources for programmatic annotation of regulatory variants and their respective target gene(s) are still lacking. Ideally, such resources would include a catalogue of regulatory regions and functional elements together with a set of impact prediction scores (40, 44). However, the resources currently available in this field are often presented in a format not suitable to this task, and information about controlled gene(s) and tissue(s) of activity is difficult to access programmatically.

To fill this gap, we have created a comprehensive resource for regulatory variant annotation including a collection of ∼2.4M regulatory elements, GREEN-DB (Genomic Regulatory Elements ENcyclopedia Database); additional functional elements (TFBS, DNase peaks, ultra-conserved non-coding elements (UCNE), topologically associating domains (TADs) and super-enhancers) and pre-processed prediction scores. Information on the controlled gene(s), tissue(s), and associated phenotype(s) are provided in GREEN-DB when possible. Here, we present a unified framework that can leverage this new resource to process standard variant call format (VCF) files and generate a comprehensive annotation of non-coding variants. We anticipate that this will aid annotation of regulatory non-coding variants identified from WGS thereby improving variant interpretation and diagnostic yield.

## Materials and Methods

### Data collection

To compile an up-to-date, comprehensive collection of regulatory elements in the human genome (GREEN-DB) we collected and aggregated information from 16 different sources, including 7 previously published curated databases, 6 experimental datasets from recently published articles, and predicted regulatory regions from 3 different algorithms. Five additional datasets were included to integrate region to gene/phenotype relationships. The full list of data sources and references is reported in Supplementary Table 1. We also collected additional data useful in evaluating the regulatory role of genomic regions, including TFBS, DNase peaks, ultraconserved non-coding elements (UCNE), super-enhancer definitions, and enhancer LoF tolerance (Supplementary Table 2) as well as 28 scores developed to predict the regulatory impact of non-coding variants (Supplementary Table 3).

### Data processing

Original tables from the various data sources were processed to generate a normalized representation of potential regulatory regions organized in a SQlite database containing information on their controlled gene(s), method(s) of detection, tissue(s) of activity, and associated phenotype(s). When needed, region coordinates were converted from GRCh37 to GRCh38 coordinates using the UCSC LiftOver tool. Additional datasets such as TAD domains, TFBS, DNase clusters, super-enhancer, UCNE, and enhancer LoF tolerance were also included in GREEN-DB with information on their overlap with the regulatory regions. The detailed description of data processing steps and SQlite database organization is provided in Supplementary Methods.

### Evaluation of collected regulatory regions

Given that creation of the GREEN-DB regions table involved a complex pre-processing of the original data sources, we first verified if these regions can still capture functional and conservation signals, a feature often reported to support regulatory role in the original data sources. First, we used Fisher’s exact test to evaluate over-representation of functional genome elements in GREEN-DB regions considering ENCODE TFBS, ENCODE DNase hyper-sensitivity clusters, UCNE regions, and a curated set of non-coding disease-associated variants (from (36)). Then, we also investigated the overlap with genomic low-complexity regions (as defined in (45)) and segmental duplications, which are mostly uninformative for variant detection. Finally, we generated a set of control regions by randomly picking from each chromosome (excluding centromeric and telomeric regions) the same number of regions seen in GREEN-DB, with comparable size distribution (Supplementary Figure 1), and compared the degree of conservation and the prediction scores distribution between these regions and the GREEN-DB regions. Using the Mann–Whitney U test we compared the fraction of bases having a PhyloP100 score above 1, 1.5, and 2 (higher values indicate more conservation) and the median and maximum score values for ncER, FATHMM MKL and ReMM scores between the GREEN-DB and control regions.

We also performed an in-depth analysis of the gene-region connections collected in the database, evaluating the distance between a region and its controlled genes, the occurrence of connections within TAD domains and the specificity of gene-region relationships (see Supplementary Methods).

### Identification of regions under variation constraint

We used data from gnomAD v3 to evaluate the possible variation constraint across GREEN-DB regions by evaluating the deviation of observed number of variants from the expectation based on a linear regression model including region length, GC percentage and overlaps with segmental duplications, low-complexity regions and exonic regions. Regions above the 99th percentiles of the resulting constraint value were considered as constrained regions. We characterised these regions and the controlled genes by looking at: (i) possible enrichment of true positive variants in the curated set of disease-causing non-coding variants from (36), Gene Ontology groups, canonical pathways, essential genes, and ClinVar pathogenic genes; (ii) distribution of the highest constraint value for each gene present in GREEN-DB; (iii) maximum constraint value between regions controlling genes in the ClinVar pathogenic or essential genes groups and all other regions in GREEN-DB. More details are described in Supplementary Methods.

### Evaluation of non-coding impact prediction scores

With the aim of providing a framework useful for variant prioritization, we reviewed the potential of 28 non-coding variant impact prediction scores to inform the analysis of WGS data from patients with rare diseases. Among these, we excluded: 10 scores not providing pre-computed values, making them difficult to apply programmatically, 2 scores developed specifically for somatic variants, and 1 score providing only disease-specific predictions for a limited set of phenotypes (see Supplementary Table 3). We used the ROCR package (46) to compare the classification performances of the remaining 19 scores when applied to a set of curated disease-causing non-coding variants from (36) including 725 true positive examples and 7,250 negative examples. More details are given in Supplementary Methods.

### A framework for regulatory variant annotation

Combining information in GREEN-DB with the three best prediction scores, population allele frequency and functional elements, we created a prioritization strategy that ranks variants overlapping GREEN-DB regions from Levels 1 to 4 providing cumulative evidence for a possible regulatory impact as described in Table 1. To further assist in result interpretation, the resulting candidate genes can be ranked based on HPO profiles using GADO (47) and the genes above 90th percentile in the ranking are suggested as likely disease related. We applied this strategy to analyze 53 WGS trios with recessive mode of inheritance and computed the resulting number of recessive and compound heterozygotes candidate variants when considering only coding variants, only GREEN-DB annotated variants or the combination of both. Details on the WGS cohort and the filtering strategy are given in Supplementary Methods.

**Table 1.** Criteria used in the prioritization strategy implemented in GREEN-VARAN annotation for non-coding variants. This is based on a four level prioritization of variants, complemented by HPO based ranking of candidate genes.

| Level | Criteria |
| --- | --- |
| Level 1 | Rare variant (population AF < 1%) overlapping one of GREEN-DB regions |
| Level 2 | Level 1 + overlap with at least one functional element among transcription factors binding sites (TFBS), DNase peaks, ultra-conserved elements (UCNE) |
| Level 3 | Level 2 + prediction score value above the suggested FDR50 threshold for at least one among ncER, FATHMM-MKL, ReMM |
| Level 4 | Level 3 + region constraint value $\geq 0.7$ |
| HPO-based prioritization | Level 4 + linked to a gene in the top 90th percentile of HPO-based ranking according to GADO prediction |

### Evaluation of validated disease-causing non-coding variants

To evaluate the performance of the proposed prioritization method, we applied it to a independent set of 45 rare disease-causing non-coding variants described in (48) (Supplementary Table 4). These variants were selected since they are independent from those used in most of the considered prediction scores. First, we computed the number of variants captured at each of the 4 prioritization levels. Then, we inserted each validated variant into WGS variants obtained for the reference sample NA12878 and evaluated the number of possible candidate variants resulting at each level assuming the disease-causing gene was known. Finally, to better assess the impact of the proposed method in reducing the number of possible candidates, we computed the number of candidate variants resulting from applying it to the whole set of WGS variants, using GADO to prioritize disease-related genes. We repeated this analysis assuming either a recessive or dominant mode of inheritance. To better evaluate how GREEN-DB annotations can help identifying regulatory variants in distant control elements, we also applied our annotations to a set of 18 previously published variants involved in human diseases and located in distant enhancers with a validated effect on gene expression (Supplementary Table 5). More details are given in Supplementary Methods.

## Results

### *The* GREEN-DB *database*

We have created a comprehensive collection of potential regulatory regions including ∼2.4M regions from 16 data sources covering ∼1.5Gb in the human genome (Figure 1A, B, and Supplementary Figure 2). A summary of the regions present in GREEN-DB is given in Table 2 with more details in Supplementary Table 6. Overall, these regions cover ∼60% of introns and ∼40% of intergenic space (Figure 1B), but overlap was observed also with UTR and other exonic regions (Supplementary Figure 3 and Supplementary Tables 7, 8). We grouped regulatory regions into five categories: bivalent (regions showing both activation and repression activity), enhancer, insulator, promoter, silencer; with enhancer and promoters representing the majority of regions (Figure 1C) with size distributions shown in Figure 1D. Each region is described by its genomic location, region type, method(s) of detection, data source and closest gene / TSS and ∼35% of regions are annotated with controlled gene(s), ∼40% with tissue(s) of activity, and ∼14% have associated phenotype(s) (Figure 1E). These data are organized in an SQLite database allowing for rapid querying based on genomic interval(s) and/or gene(s) of interest (the database structure is described in Supplementary Results and in Supplementary Figure 4). GREEN-DB regions are also provided as extended BED files for integration into existing analysis pipelines.

**Table 2.** Summary of GREEN-DB information, reporting regions counts and number of genomic bases covered.

| GREEN-DB | GRCh37 |  |  | GRCh38 |  |  |
| --- | --- | --- | --- | --- | --- | --- |
|  | No. of Elements | Mean size (bp) | Bases covered | No. of Elements | Mean size (bp) | Bases covered |
| Enhancer | 1,834,183 | 1107 | 1,450,755,698 | 1,832,830 | 1,107 | 1,449,153,178 |
| Promoter | 566,102 | 573 | 234,890,654 | 565,323 | 573 | 234,315,553 |
| Silencer | 4,306 | 208 | 895,868 | 4,302 | 208 | 894,792 |
| Bivalent | 8,413 | 1,333 | 11,215,000 | 8,409 | 1,333 | 11,210,309 |
| Insulator | 23 | 846 | 17,504 | 23 | 846 | 17,504 |
| All regions | 2,413,027 | 981 | 1,504,116,499 | 2,410,887 | 981 | 1,502,180,018 |

**Figure 1.**
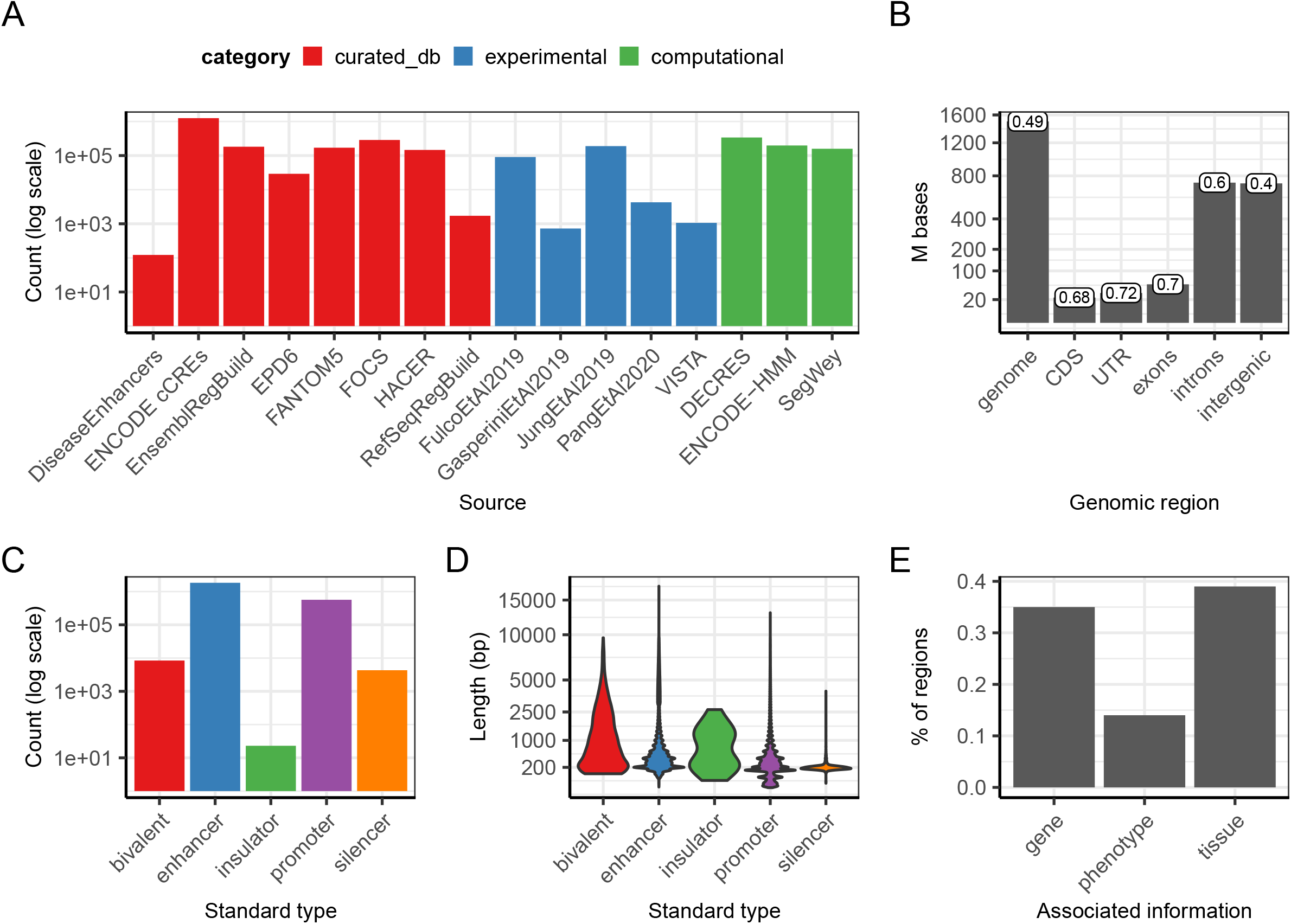
Summary statistic of regions collected in the GREEN-DB. (A) GREEN-DB collects human regulatory regions from 16 different sources including curated databases, experimental assays, and computational predictions. (B) Number of bases captured by these regions across different genomic locations and covered fraction of each genomic location (label on top of bars). (C) GREEN-DB contains bivalent, enhancer, insulator, promoter, and silencer regions with sizes mostly between 100 and 1000 bp (D). (E) Fraction of regions with associated gene, phenotype and tissue information. Phenotype information was derived from GWAS studies (via overlap of significant SNPs with GREEN-DB regions), HPO (via controlled genes), and DiseaseEnhancer dataset.

Processed regions in GREEN-DB maintain strong support for a regulatory role as indicated by the enrichment of several functional genomic signals including transcription factor binding sites (TFBS, OR 9.67), DNase hypersensitivity peaks (OR 13.13), GTeX significant eQTLs (OR 2.93), ultra-conserved non-coding elements (UCNE, OR 8.35) and a curated set of non-coding disease-causing mutations (OR 2.05). At the same time, they are depleted for difficult-to-address regions such as segmental duplication (SegDup, OR 0.45) and low-complexity regions (LCR, OR 0.22) (Supplementary Figure 5 and Supplementary Table 9). Compared to random regions with comparable size and distribution across the genome, GREEN-DB regions showed a larger proportion of bases with PhyloP100 score above 1, 1.5, and 2 (p-value < 2.2E-16 for all comparisons, Mann–Whitney U test) as well as higher per-region median and maximum values for ncER, FATHMM MKL and ReMM scores (p-value < 2.2E-16, Mann–Whitney U test) (Supplementary Figure 5).

### Controlled genes annotation in GREEN-DB

Overall, ∼88% of GREEN-DB regions have a putative association to one or multiple genes, either because these associations were determined experimentally (∼32%), or because of a gene in close proximity (distance ≤ 10kb, ∼56%) (Figure 2). Considering the 837,879 regions with a validated region-gene association, they interact with a total of 48,230 different genes, covering 67% of all genes and 97% of protein-coding genes from ENCODE v33 basic set. Controlled genes also cover 97, 98, and 100% of clinically relevant genes from PanelApp (49), ClinVar (pathogenic genes only), and ACMG actionable genes list, respectively (Supplementary Table 10). Considering controlled genes, we saw that the closest gene is among annotated controlled genes only for ∼70% of enhancers and ∼25% of silencers, while this proportion is much higher (∼90%) for promoters, as expected. The closest gene is not the only one controlled in 10-25% of cases for distal control elements, even when the element overlaps a specific gene (Figure 2D). The region-to-gene relationships showed a high degree of specificity, with most regions controlling less than 5 genes, while most genes are controlled by multiple regions (Supplementary Figure 6 and 7). Additional details on the genes’ regulatory space are provided in Supplementary Results.

**Figure 2.**
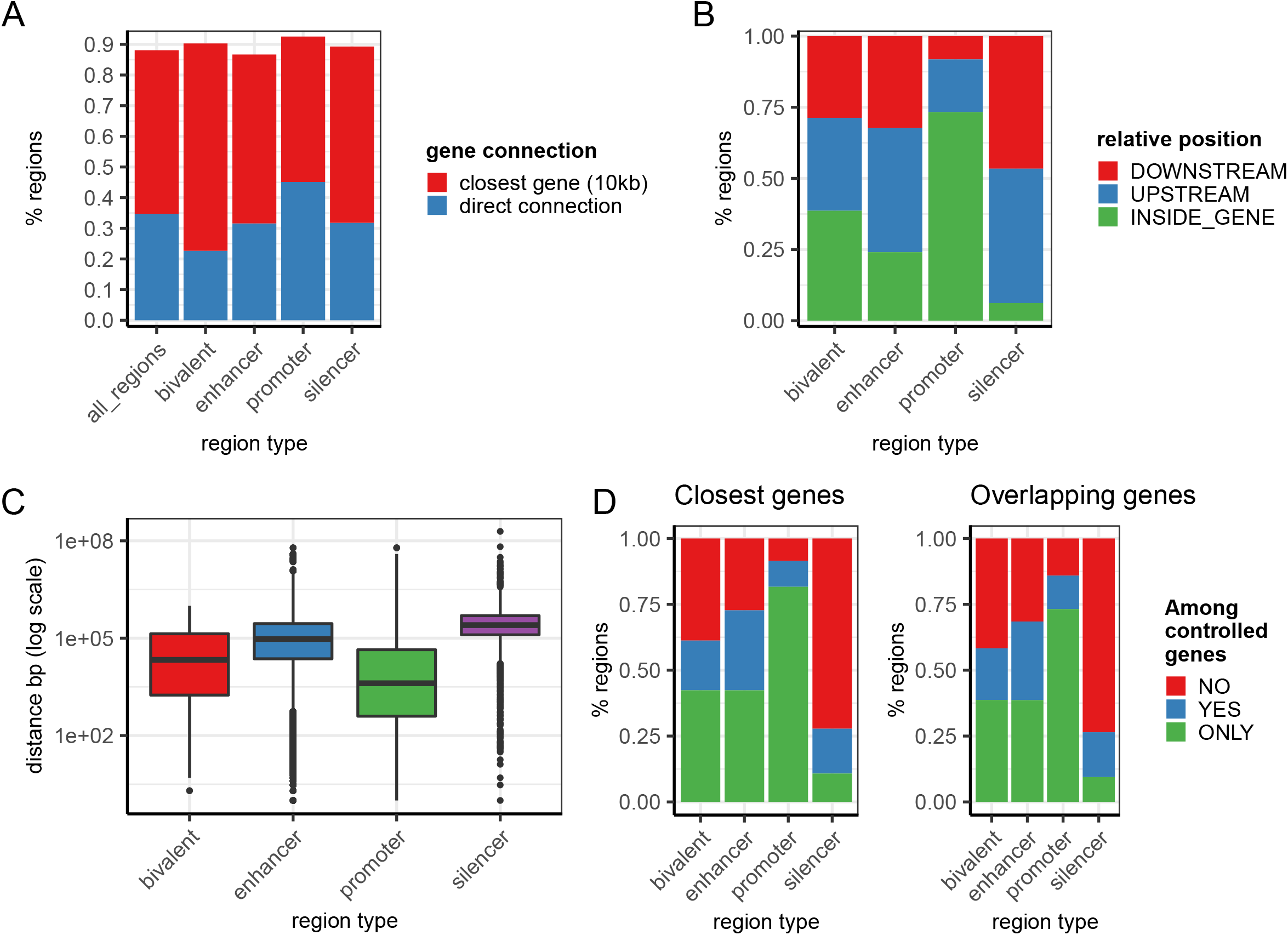
Gene regulatory space. (A) Overall 839,807 regions (∼35%) in GREEN-DB are experimentally associated with a controlled gene, and the fraction of regions with a plausible associated gene reaches ∼90% when we include close genes within 10kb distance. (B) Considering only experimental associations, distant control elements are mostly located up- or downstream of a gene, with a smaller proportion observed within genes. The large proportion of gene overlap observed for promoters is mostly explained by their partial overlap with the first gene exon. (C) The distance between a region and its controlled gene(s) is larger for enhancers, silencers, and bivalent, with most regions located between 10kb and several Mb away from the controlled gene. (D) Interestingly, a large proportion of these regions may not control their closest gene(s) even when they are located within a specific transcript.

### Regions constrained against sequence variation

We calculated a constraint metric for GREEN-DB regions ranging from 0 to 1, so that regions with higher values have lower than expected numbers of variants. Based on this metric we defined as constrained the 23,102 regions above the 99th percentile of the distribution (mostly enhancers and promoters, Supplementary Figure 8). Comparison with other regions in GREEN-DB showed that constrained regions are more conserved (Supplementary Figure 8C) and enriched for tissue or gene-specific regions (p < 2.2E-16, Supplementary Figure 9) and for true disease-causing mutations from the curated set (OR 13.32, 95CI 8.32-21.55, p 1.42E-27).

Overall, constrained regions control 4,579 genes and these genes are strongly enriched for essential genes and genes bearing pathogenic variants in ClinVar (FDR 4.53E-133 and 6.21E-198, respectively). Complete enrichment results are reported in Supplementary Table 11. When comparing the maximum constraint value of associated GREEN-DB regions, genes in the ClinVar pathogenic and essential groups are controlled by regions with higher constraint value compared to other genes (p < 2.2E-16, Matt-Withney U test, Supplementary figure 10). Overall, regions above the 70th percentile control > 95% of ClinVar pathogenic genes, essential genes and are associated with genes with lower observed/expected ratio for loss of function variants in GnomAD (oe_lof) (Figure 3). However, the median constraint value considering all associated regions is only slightly higher for pathogenic and essential genes compared to normal genes (Supplementary Figure 10).

**Figure 3.**
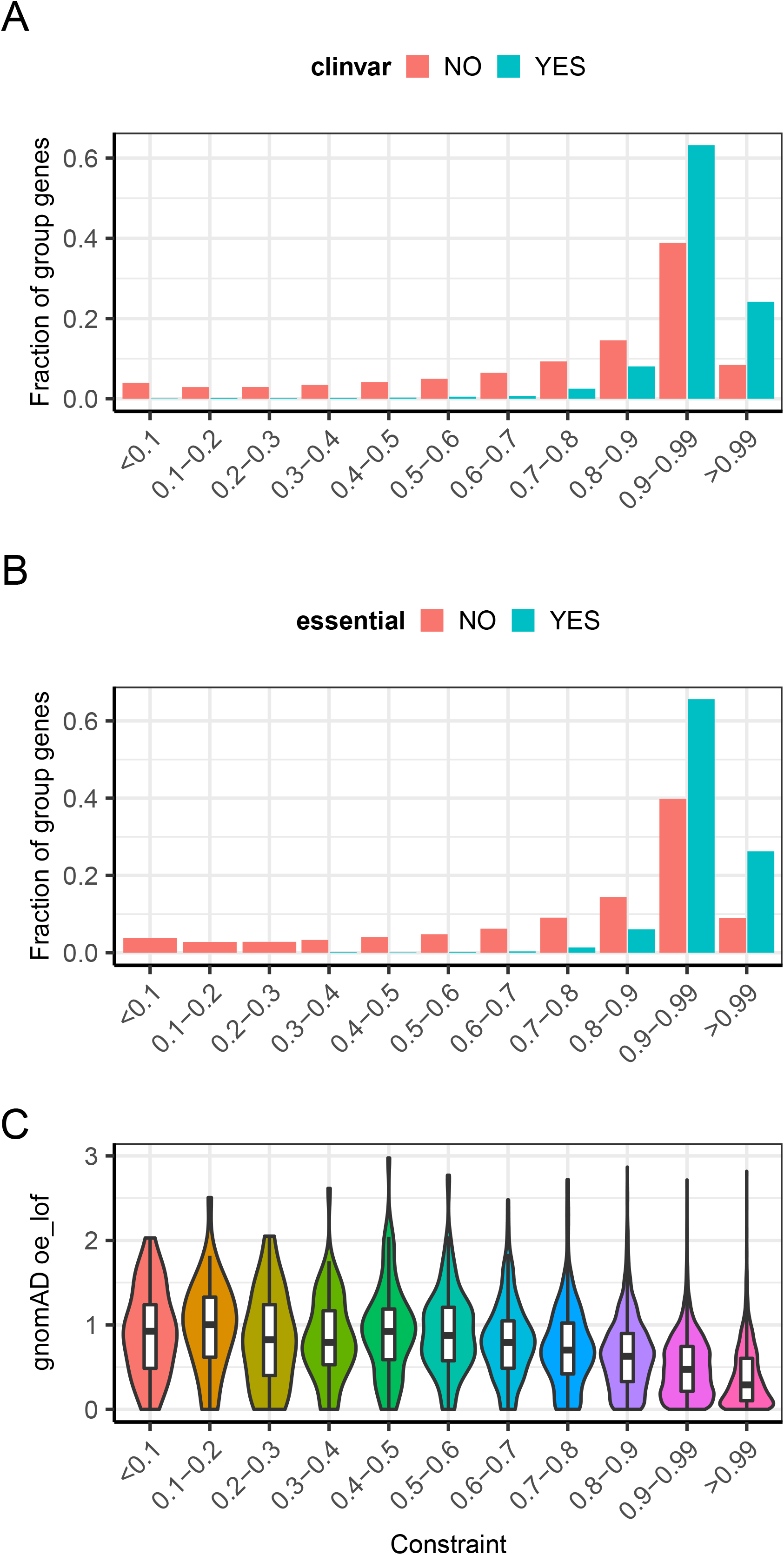
Constraint regions control diseases-associated and essential genes. For various constraint value tranches, we calculated the fraction of ClinVar (A) or essential (B) genes from mouse knockout screens controlled by at least one region in the corresponding tranche. Both groups show a large fraction of genes controlled by regions with constraint value ≥ 0.9 and > 95 % of genes in each group are linked to a region with constraint ≥ 0.7. Regions with high constraint are also controlling genes with lower oe_lof value in gnomAD (C), suggesting they are associated with genes intolerant to variations.

### Evaluation of non-coding impact prediction scores

We considered 28 previously published prediction scores that can be applied to evaluate the impact of non-coding variants. Of these, 13 do not provide pre-computed values or were developed for somatic variants only and were thus removed from further analyses, while GWAVA and EIGEN provide 3 and 2 possible values respectively. Using a curated set of disease-causing, non-coding variants from (36), we evaluated the performance of 18 scores in classifying disease-causing variants. The FINSURF algorithm obtained the best overall classification result (OPM 0.73, AUC 0.94), but the pre-computed scores only cover 15% of the genome, limiting its application in WGS annotation. We selected ncER, FATHMM-MKL and ReMM as the best scores combination that provided both high genomic coverage (> 90%) and good classification performances (OPM > 0.4 and AUC > 0.8) (Supplementary Table 12). Overall, no single score seemed able to robustly remove false-positive calls while maintaining high sensitivity. Indeed, when TPR is set to 0.9, the FDR is above 0.5 for all scores, while controlling the FDR ≤ 0.5 results in TPR values below 0.5 for most scores. To assist the use of these scores in variant analysis, we also computed the score thresholds corresponding to TPR ≥ 0.9, FDR ≤ 0.5, and maximum accuracy (detailed metrics are shown in Supplementary Figure 12 and Supplementary Table 13).

### A framework for annotation and prioritization of non-coding variants from WGS

We created a tool (GREEN-VARAN: Genomic Regulatory Elements ENcyclopedia VARiant ANnotation) to annotate VCF files with information from GREEN-DB. The tool is written in the Nim programming language using hts-nim (50) and processes standard VCF files by adding annotation on overlapping regulatory region(s) type(s), IDs and constraint values, controlled gene(s) and closest gene(s) with their distance. The tool can also update existing gene consequence annotations from snpEff or bcftools and a tag can be added to highlight variants linked to gene(s) of interest. When allele frequency, non-coding prediction scores and functional element annotations are present, GREEN-VARAN also classifies variants according to the 4 levels described in Methods. However, the prioritization strategy is fully configurable to be able to take into account additional custom annotations present in the input VCF file. Additional pre-processed datasets useful for annotation and a Nextflow workflow are distributed together with GREEN-DB and can be used to generate a fully annotated VCF for non-coding variants. Given an annotated VCF, a list of variants or a list of GREEN-DB IDs, GREEN-VARAN can also be used to query GREEN-DB and retrieve detailed annotations including tissue(s) of activity, data source(s) and associated phenotype(s). More details on the tool, available datasets and the annotations added to VCF are given in Supplementary Results.

### GREEN-VARAN annotation captures validated non-coding, disease-associated variants

We used an independent set of 45 rare curated variants from (48) (Supplementary Table 4) to evaluate how GREEN-VARAN annotations can help capture disease-causing variants in non-coding regions. When applying the proposed prioritization system, we observed 40 (89%), 35 (78%), 32 (71%) and 9 (20%) validated variants captured at Level 1, 2, 3 and 4, respectively. When we added these known variants into WGS variants from the reference NA12878 sample and looked at variants associated to the relevant gene only, GREEN-VARAN prioritization was able to reduce the number of candidate variants to < 25 and < 5 for heterozygous and homozygous variants, respectively. The possible candidates were reduced to < 5 and just one when considering level 3 variants, even if 29% of causative variants were lost (Figure 4A). In a scenario where the disease gene is not known, we integrated GREEN-VARAN annotations with HPO-based prioritization using random HPO terms associated with the relevant disease for each validated variant. Using mild filtering (Level 1 variants and 0.9 GADO threshold) this approach generated 13k - 23k and 1k - 2k candidate variants for dominant and recessive inheritance respectively, while a more stringent filtering (Level 3 variants and 0.95 GADO threshold) reduced the number of candidates to 400 - 800 and 8 - 32, while still capturing 71% of the validated examples (Figure 4B, Supplementary Table 14).

**Figure 4.**
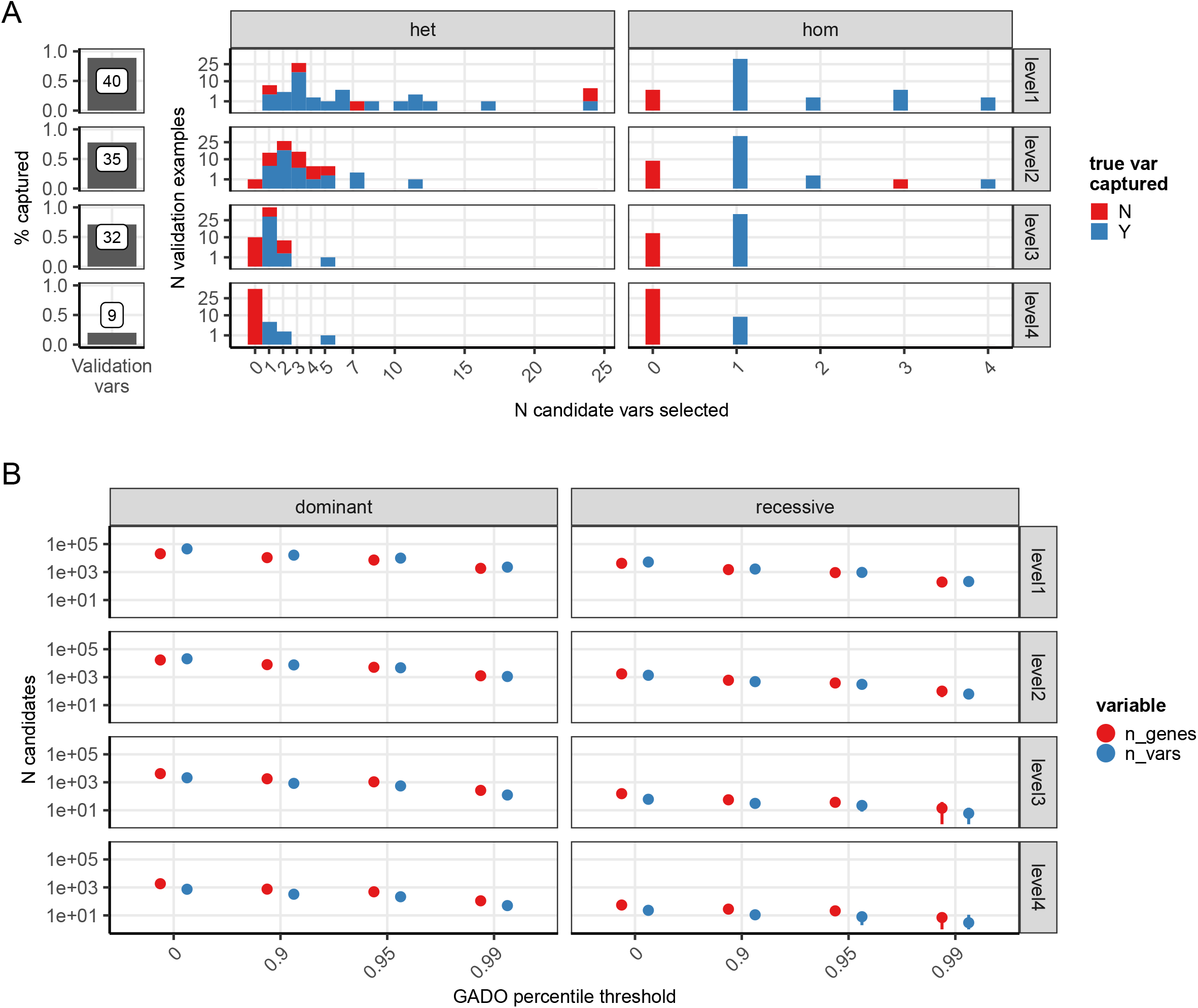
Application of GREEN-VARAN to validated variants. We inserted a set of 45 validated variants into variants from the reference sample NA12878 and tested how GREEN-VARAN annotations are able to capture them and how many candidate variants are selected when considering only the disease-causing gene (A). We then evaluated how many candidate variants / genes are selected by our prioritization method in combination with HPO based gene ranking when the disease gene is not known (B).

Most of the above validated variants in non-coding regions are actually located in UTR or intronic regions of the controlled gene; thus we extended our analysis to a set of 18 previously published variants located in distant enhancers with demonstrated regulatory effects on a disease gene. GREEN-DB annotations were able to capture all tested variants, linking them to the expected gene. When applying the proposed prioritization strategy we found 18, 13 and 2 variants ranked at Levels 2, 3 and 4, respectively. Details on each variant are reported in Supplementary Table 5.

### GREEN-VARAN annotation reveals new candidate genes in WGS trio analysis

To evaluate the impact of adding non-coding annotations in a more realistic scenario, we applied our annotation framework to the analysis of small variants from 53 non-consanguineous WGS pedigrees. The main aim was to evaluate the number and relevance of new candidate variants and genes, considering that the interpretation of VUS and novel genes is time consuming and expensive and thus it is important that new tools do not add too many false-positives. Considering variants identified in each individual, we found a median of ∼75.7k rare variants (population AF < 0.01), including ∼38k in GREEN-DB regions (Level 1), ∼17k also overlapping functional signals (Level 2) and ∼1.5k further prioritized based on prediction scores (Level 3) (Supplementary Figure 13). When considering rare recessive variants that segregate with the phenotype in each pedigree, adding GREEN-DB annotations increased the number of candidate variants from 0 - 22 (exonic variants only) to 83 - 2209 (GREEN-VARAN Level 1). Looking at variants with stronger support for regulatory impact, the number of candidates is reduced to 45 - 1291 at Level 2, 0 - 75 at Level 3 and 0 - 47 at Level 4 (Figure 5A, B). When considering compound heterozygotes involving a protein-changing variant we observed 853 - 3362 combinations with a Level 1 variant and 13 - 112 with a Level 3 variant (Figure 5C). The new candidates identified also included genes likely to be relevant to the disease phenotype based on HPO profiles. Indeed, when restricting to genes with high GADO score the number of recessive candidates is increased from 0 - 2 (exonic variants only) to 0 - 9 (Level 3 GREEN-DB variants), and we identified 0 - 10 new compound heterozygotes involving a protein-changing and a Level 3 variant. A similar trend was observed also when considering only clinically relevant genes from PanelApp or Clinvar. Detailed counts for candidates identified in the trio analysis are reported in Supplementary Table 15.

**Figure 5.**
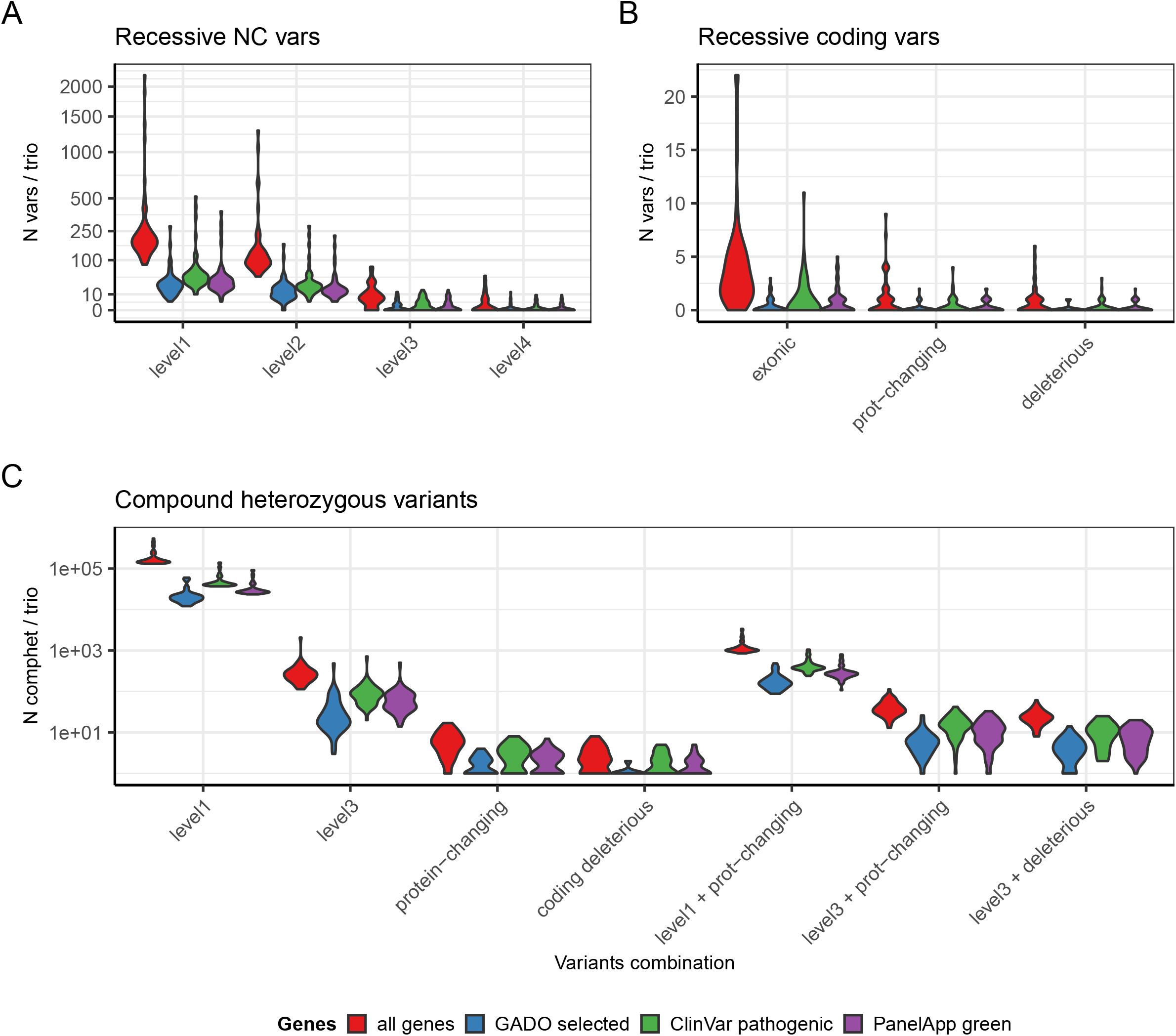
Impact of non-coding annotations on WGS variant prioritization. The violin plots represent the number of recessive candidate variants found in 53 WGS trios for non-coding variants prioritized by GREEN-VARAN (A) or considering only coding variants (B). We then considered compound heterozygotes combinations (C) reporting the number of candidates when considering only GREEN-VARAN prioritized variants (Level 1 and Level 3), only coding variants or combinations involving one coding and one non-coding variants. In all plots the counts are reported for all genes, for genes ranked above the 90th percentile in GADO distribution (GADO selected), for ClinVar pathogenic genes and for PanelApp disease genes.

## Discussion

Variants in non-coding regions of the genome have clearly been implicated in disease risk from both GWAS studies (14–16) and WGS rare disease studies (8, 10–12) and the large fraction of missed diagnoses in WES studies (51–53) indicate that the non-coding genome is likely to harbor many variants of clinical diagnostic significance and may account for low diagnostic yield in clinical WGS rare disease studies. Whilst information about types and locations of regulatory regions has previously been described in the literature (17, 19–25), the systematic interrogation of these in whole-genome sequencing data remains challenging and limited by the lack of resources to readily access these programmatically (54–56). To fill this gap, we have developed a framework for the automated annotation of WGS variants compiling an extensive catalogue of regulatory regions and developing accompanying tools and resources that can be integrated into routine bioinformatics pipelines to annotate non-coding variants and improve their interpretation and prioritization in rare disease WGS datasets.

We have collected and curated data from published, experimental, and computational sources to create a database providing a standardized representation for ∼2.4 million regulatory elements in the human genome (GREEN-DB). To support the interpretation of the impact of genetic variants, each regulatory region is annotated with a rich set of information including controlled gene(s), closest gene(s), tissue(s) of activity, potentially associated phenotype(s) from GWAS studies and Human Phenotype Ontology, and a constraint metric representing the tolerance to genetic variation. To interpret the biological role of a regulatory region, it is essential to know the genes it controls and in which tissues it is active. GREEN-DB contains experimentally validated region-gene links and tissue information for ∼35% and ∼40% of the regions, respectively. Overall, the database provides regulatory information for 48,246 genes, including most of the clinically relevant genes from PanelApp, ClinVar and ACMG, supporting its usefulness in human disease research.

Our analysis confirms the complexity of the relationship between regulatory regions and controlled genes that can not easily be explained by spatial proximity in the (linear) genome as previously demonstrated, e.g., by high-throughput studies of chromatin interactions (32, 57–59). Indeed, for silencer and enhancer elements, the controlled gene was the closest gene in only 10% and 40% of cases respectively, whilst regulatory regions within a gene exert regulatory control on that specific gene in less than 60% of the cases. Even if we cannot exclude that these observations may be influenced by incomplete annotation of controlled genes, this has considerable implications for the interpretation of non-coding variants where the search for disease-associated genes often starts from the closest genes (60, 61). Overall, we observed a high degree of specificity in the region-gene relationship, with a large fraction of regions controlling less than 5 genes, while most genes are controlled by multiple regulatory regions suggesting that multiple, spatially distant, genomic regions may have similar phenotypic impacts. A correlation emerged between the number of controlled genes and the number of active tissues for each region, confirming the tissue-specific nature of gene regulation and supporting the idea that alterations in a regulatory region can have different impacts in different tissues (12). This makes the comprehensive annotation of all regions that influence or regulate the normal activity of a gene so important for understanding the consequences of genomic variants on the respective gene function.

We also integrated information from GWAS studies and HPO databases to provide a possible associated phenotype for ∼15% of the regions. This resource will be useful for the interpretation of new variants found in the regulatory regions, providing hypotheses on their potential biological impact. The fact that only a limited number of regions has an associated phenotype, despite a large number of GWAS hits available (62–64), can partially be explained by a reduced phenotypic effect of regulatory regions alterations due to complex regulatory network and tissue-specific effects. On the other hand, this also underlines how the impact of rare disrupting variants in the non-coding space is largely unexplored and how resources like GREEN-DB can inform our understanding of human diseases.

We calculated a constraint metric that reflects the tolerance of each region to sequence variations. The maximum constraint value for regions controlling essential genes and genes involved in human diseases is significantly higher compared to other genes and constrained regions (constraint value ≥ 0.99) are associated with genes under variation constraint (low observed/expected ratio for loss of function variants in gnomAD) and strongly enriched for essential genes and genes involved in human diseases. Thus, our constraint metric can be used as a stringent filter to prioritize regions likely to be relevant in controlling disease genes, even if the redundancy of the regulatory network implies that also more variable regions may be involved as suggested by the analysis of median constraint across all regions associated with ClinVar pathogenic genes.

To further assist the interpretation of variants located in regulatory regions, we collected pre-computed values from 19 different impact prediction algorithms and compared their ability to classify a curated set of established disease-causing non-coding variants. Overall, we must take into account that such comparisons are limited by (i) the nature of the known variants collected so far, which are mostly variants within the affected gene and poorly capture distant regulatory elements (56); and (ii) the potential overlap of the test variants with the training sets used by each algorithm, which are often unknown. Based on classification performances and genome coverage, ncER (65), FATHMM (66, 67) and ReMM (38) algorithms emerged as the best performing scores, probably reflecting their specific training on disease-associated variants.

We developed a tool (GREEN-VARAN) that integrates information from GREEN-DB, non-coding impact prediction scores, functional elements and population AF annotations, into a 4 level prioritization system to rank the regulatory potential of non-coding variants in a VCF file. The proposed approach can effectively capture variants involved in human diseases, as shown by our ability to recapitulate known disease-associated variants from the literature.

When applied to a set of 45 validated non-coding variants, our approach associated 40 of them to the correct gene and classified 32 as likely impacting gene expression (prioritization level 3), while considerably reducing the number of candidate variants to be evaluated for the causative gene (in 25 cases the causative variant was the only one selected). When the disease gene is unknown, the integration of GREEN-VARAN prioritization with HPO-based gene ranking, greatly reduces the number of candidates to evaluate in a WGS singleton even if the number of candidates remains challenging (up to 1500 genes) in case of the dominant inheritance model.

Since the considered validated variants were mostly located within the controlled gene, we further tested GREEN-VARAN prioritization on 18 variants in distant enhancers from (68–70) confirming its ability to identify the proper controlled gene and prioritize relevant variants (13 out of 18 were classified in Level 3 as likely impacting gene expression).

Application of our new annotation system to a dataset of 53 WGS trios highlighted its potential impact for rare-disease variant analysis. When considering rare recessive variants that segregate with the phenotype in each pedigree, the number of candidate variants is greatly increased by the addition of GREEN-VARAN annotations. However, when considering only variants with stronger support for regulatory impact, the number of candidates is reduced to an amount manageable for downstream analysis (0 - 75 at Level 3 and 0 - 47 at Level 4, Figure 4A, B). Whilst this number of candidates is still too many for a diagnostic lab to consider, this is certainly in the realms of the possible for research-based inspection, especially in otherwise difficult-to-solve cases. The new variants prioritized by GREEN-VARAN are of particular interest in compound heterozygote candidates, especially when non-coding variants create new combinations with a prioritized coding variant, an approach that already resulted in increased diagnosis in a recent large clinical WGS study (71). Interestingly, we observed 13 - 112 combinations with a Level 3 variant and the new candidates identified included genes likely relevant for the disease phenotype based on HPO profiles. A similar trend was observed when considering clinically relevant genes from PanelApp or Clinvar suggesting that inclusion of our non-coding annotation can reveal previously ignored candidate genes likely to have an impact on patient phenotype.

In summary therefore, we have developed a highly curated dataset of regulatory regions (GREEN-DB) and integrated it with functional elements and prediction scores into a new framework for the annotation and prioritization of regulatory variants in WGS analysis. We provide a complete annotation workflow implemented in Nextflow that uses our GREEN-VARAN tool to prioritize regulatory variants in VCF files from whole genome sequencing data. The resources presented here therefore represent a significant advance for researchers engaged in analyzing patient genomes and will be useful for rare disease research as well as for the interpretation of common disease variants from GWAS.

## Data availability

GREEN-DB BED files and SQlite database are available from https://zenodo.org/record/3981033. GREEN-VARAN tool, the annotation workflow and additional data are available from https://github.com/edg1983/GREEN-VARAN.

## Acknowledgments

Authors acknowledge Dr. Dimitris Vavoulis for critical discussion on the constraint model analysis.

## Additional files

Additional File 1. Supplementary Tables 1-15 (.xls)

Additional File 2. Supplementary Methods and Results. Supplementary Figures 1-14 (.pdf)

