## Additional file 2 - Supp methods and figures for "GREEN-DB: A framework for the annotation and prioritization of non-coding regulatory variants from whole-genome sequencing data"

Giacopuzzi E
Popitsch N
Taylor JC

**Index**

| | |
|---|---|
| **Supplementary Methods** | **2** |
| Data processing | 2 |
| Pre-processing of scores predictions and additional datasets | 4 |
| Genes lists used in this study | 4 |
| Analysis of gene regulatory space | 5 |
| Identification of regions under variation constraint | 6 |
| Evaluation of non-coding impact prediction scores | 6 |
| Prioritization strategy | 7 |
| Evaluation on validated disease-causing non-coding variants | 7 |
| Detection of variants in WGS trios | 8 |
| **Supplementary Results** | **9** |
| GREEN-DB Database structure | 9 |
| Controlled genes annotation in GREEN-DB | 10 |
| Pre-processed datasets available in the GREEN-DB collection | 10 |
| The GREEN-VARAN tool and annotations | 11 |
| Nextflow annotation workflow | 12 |
| **Supplementary References** | **13** |
| **Supplementary Figures** | **14** |

**Supplementary Methods**

*Data processing*

All data in GREEN-DB was processed to generate a standardized representation of regulatory regions, their controlled gene(s), method(s) of detection, and tissue(s) of activity. First, we collected a set of possible regulatory elements from 16 different sources including recently published datasets and previously assembled curated databases (Supplementary Table 1). For 3 computational datasets, only regions predicted as active were retained:

- DECRES: predicted active-promoters (A-P) and active-enhancers (A-E) from supplementary table 2 in (Li *et al.*, 2018)
- ENCODE-HMM: the predictions on regulatory region activities across 9 cell lines were downloaded from UCSC genome browser and all regions classified at level 8 or below were selected (active promoter, weak promoter, poised promoter, strong enhancer, weak enhancer, insulator).
- SegWey: predictions of regulatory regions activities across 164 cell types were downloaded from SegWey website. Based on the description given in the original paper we selected only high confidence active regions, identified by the labels "Promoter", "Enhancer", "Bivalent".

Genomic coordinates were then parsed from each raw dataset, and each region was assigned a standard region type (bivalent, insulator, promoter, enhancer, silencer) and a unique identifier, used to link the region with all the additional annotations. This standard region type was assigned based on the definition reported in the source dataset after removing detailed classifications such as weak, strong, etc. Additional information reported about tissues of activity, controlled genes, methods of detection, and associated phenotypes was extracted into 4 separate tables: tissues, genes, methods, and phenotypes. The standardized region tables from each dataset were then concatenated in a single dataset containing 3,107,738 regions, covering ~49% of the genome. To maximize the informativeness of the final collection and remove potential spurious data, we applied several processing steps:

1. CAGE-seq results are represented by multiple small fragments of a few hundred bases that collectively map to gene promoters. Thus, to obtain a less redundant dataset of possible promoters, we collapsed to single intervals CAGE-seq peaks within 500 bp from each other. When associated transcript information was available, we collapsed peaks only if they were associated with the same transcript.
2. Some techniques such as HiC could have a poor resolution or generate spurious associations, so we analyzed the distribution of regions sizes and number of regulated genes per region and removed extreme outliers. Specifically, we removed from the region and gene tables, regions with dimension above the 99th percentile and region-gene interactions for regions with a number of associated genes above the 99th percentile;
3. The various data sources used to compile our database partially rely on the same set of experimental data and thus some regions may be represented multiple times. Thus,

to reduce redundancy, regions with the same standard type and a reciprocal overlap ≥ 50% were collapsed into single entities;

4. Additionally, controlled genes were added for each region based on overlap with significant GTeX eQTLs. For each region, we added as a potential controlled gene any gene having a significant eQTL located within the region;
5. When possible, aliases and old gene symbols were converted to official gene symbols from HGNC and the corresponding Ensembl gene ID was added;
6. Interactions reported in the genes table were compared with TAD domains definitions from the TAD KB database and annotated as occurring in the same TAD or not.
7. Associated phenotypes were added for each region based on the overlap with genome-wide significant SNPs from 2 catalogs of GWAS studies (GRASP and GWAS catalogue). For each region, we added as associated phenotypes all traits having a significantly associated SNP ($p < 5E-08$) located within the region;
8. Additional phenotypes were added from Human Genotype Ontology (HPO) based on the controlled genes. For each region, we added as associated phenotype all HPO terms annotated for the controlled genes;
9. Since most of the original data sources were available only for hg19/GRCh37 assembly, the final region table was converted to GRCh38 by coordinates lift-over using UCSC LiftOver tool;
10. As an additional sanity check, any gene-to-region connection for which the gene and region were located on different chromosomes were removed as likely artifacts derived from erroneous gene id assignment or coordinates lift over process.
11. Each region was also annotated with the closest gene and closed TSS for all genes or only protein-coding genes based on the GENCODE v33 basic set;
12. Each region was annotated with the percentage of bases having a PhyloP100 value above 2, 1.5, or 1 as well as the median and max PhyloP100 value across the region.

Processed data were then compiled in an SQLite database with 6 tables describing the GRCh37 / GRCh38 regions and their associated genes, tissues, phenotypes, and methods of detection, linked by the unique region IDs. Information on the original data source from which the record is derived is included in each table as well.
Similarly, we collected 6 additional datasets useful for annotation of regulatory regions: Topologically associating domains (TAD), transcription-factor binding sites (TFBS), DNase hypersensitivity clusters, super-enhancers, ultra-conserved noncoding elements (UCNE), and enhancer LoF tolerance probability (Supplementary Table 2). All these regions were assigned a unique identifier and converted to GRCh38 coordinates using UCSC LiftOver when necessary. When available, information about tissues of activity was stored in a dataset-specific tissue table. These processed tables were incorporated into the SQLite database and the overlaps between the additional regions and GREEN-DB regions were pre-computed and stored in ID-to-ID link tables to allow the rapid collection of all available information for each region described in GREEN-DB.

For both GRCh37 and GRCh38 GREEN-DB regions, we also generated a BED-like table with chromosome, start and end coordinates, and region ID in the first 4 columns and including additional information useful for variant annotation like region type, constraint metric, number of supporting methods, phyloP100 median conservation value, closest gene/TSS with distance, and controlled genes.

*Pre-processing of scores predictions and additional datasets*
We pre-processed all the additional datasets described in Supplementary Table 2 and the pre-computed values for scores in Supplementary Table 11 to create a set of standard tables in BED-like format. For each dataset we parsed information from the original tables to have genomic coordinates in the first 3 columns and a relevant region ID in column 4, which can be either a unique region ID or the name of the binding TF for TFBS. All prediction scores pre-computed values formatted in table format reporting genomic coordinates, reference and alternate alleles in columns 4 and 5 when available, and then the score value. All tables were then binary compressed and indexed for easier integration into traditional bioinformatics pipelines.

*Genes lists used in this study*
For Gene Ontology groups, canonical pathways, and REACTOME pathways we used gene symbols lists from MSigDB c5 and c2 collections v7.1 downloaded from http://www.gsea-msigdb.org/. Essential genes lists were obtained from https://github.com/macarthur-lab/gene_lists (core_essential 283 genes; mgi_essential 2,454 genes). Clinvar pathogenic genes were extracted from ClinVar variant summary data (https://ftp.ncbi.nlm.nih.gov/pub/clinvar/tab_delimited/) accessed on 15/10/2021, by selecting only genes reported as pathogenic/likely pathogenic with no conflicting interpretations (4,967 genes). PanelApp green genes were obtained from 321 disease gene panels downloaded from PanelApp (https://panelapp.genomicsengland.co.uk/) on 04/05/2020.

*Curated variant sets used in this study*
In this study we used different sets of curated disease-causing non-coding variants from previous studies to either evaluate the performance of impact prediction scores or the ability of GREEN-VARAN annotation to prioritize disease-causing variants in the non-coding space. A summary of the curated sets used is reported in the following table

| Source publication (PMID) | N variants | Section where described | Application in this manuscript | Description |
|---|---|---|---|---|
| 30744685 | 7,975 (725 true positive examples; 7,250 negative | Supplementary Tables 1, 4 and 5 from source publication | Evaluation of non-coding impact prediction scores | A manually curated collection of disease-causing non-coding variants collected from ClinVar and HGMD and an associated set of “region-context” matched negative examples |

| | | | | |
|---|---|---|---|---|
| | examples) | | | |
| doi.org/10.1101/2021.05.03.442347 | 49<br>(45 rare) | Supplementary Table 4 in this paper | Evaluation of validated disease-causing non-coding variants | These established disease-causing variants were selected since they are independent from those used in most of the considered prediction scores. Thus they represent an unbiased set to test the ability of our prioritization approach in capturing disease relevant variants. |
| 31395865<br>24357527<br>doi.org/10.1101/2020.02.21.959734 | 18 | Supplementary Table 5 in this paper | Evaluation of validated disease-causing non-coding variants | A collection of previously published disease-associated variants located in distant enhancers. Since the validated examples from the other datasets are mostly represented by variants located within a gene, we wanted to specifically test our annotations against a set of variants in distant enhancers. |

*Analysis of gene regulatory space*

We evaluated the relationship between regions and associated genes for the 837,879 regions for which we have collected experimentally supported associations. Based on gene definitions from the GENCODE v33 basic set, we investigated where each region was located with respect to each of its associated genes (upstream, downstream, or overlapping the gene) and the region-gene distances for each of the 4 main region types having associated genes (bivalent, enhancer, promoter, silencer). Finally, we evaluated the proportion of regions for which the closest gene/TSS was among or the only controlled gene, and the proportion of regions located within a gene, but controlling other distant ones. For the 48,246 genes present in the GREEN-DB annotations, we calculated the number of associated genes per region (GxR) and the number of associated regions per gene (RxG). We correlated the GxR value with the number of tissues per region to assess whether regions controlling multiple genes are more likely to do so in a tissue-specific manner. Correlation significance was tested using Spearman's correlation test. We then selected 491 genes with an extremely large regulatory space, defined as those in the 99th percentile of RxG distribution (genes with at least 182 associated regions), and used the hypergeometric test to assess their enrichment across Gene Ontology groups and canonical pathways from MSigDB v7.1 as well as essential genes derived from cell-culture or mouse knock-outs and genes bearing any pathogenic/likely pathogenic mutation in ClinVar. FDR of the 13,960 performed tests was controlled using the Benjamini–Yekutieli method (Benjamini and Yekutieli, 2001).

*Identification of regions under variation constraint*

To evaluate the possible variation constraint across GREEN-DB regions, we computed for each region the number of overlapping PASS variants from gnomAD v3 WGS dataset. To prevent the detection of false-positive constrained regions due to partial inaccessibility, we filtered out regulatory regions that overlap more than 50% with known segmental duplications (segdup) or low-complexity regions (LCR). We furthermore removed regions on chrY and chrM leaving us with 2,310,114 regulatory regions. For these regions, we computed the region's GC density (GC) as a proxy for the region's mutability owing to the spontaneous deamination of methylated cytosines, and the fraction of bases overlapping gene exons (exonic), to compensate for higher sequence constraint in coding regions. We then created a linear regression model with the number of variants as the dependent variable:

$$N_{var} = length + GC + segdup + LCR + exonic$$

$N_{var}$, GC density, sequence length, and exonic variables were transformed to approximate normality using Blom's transformation (Blom, 1958). In the case of LCR and segdup, the majority of values were equal to 0 making the above transformation ineffective, so we treated these two variables as binary by setting all non-zero values equal to 1. The residuals from the model were ranked from lowest to highest, and assigned a percentile such that regions with the lowest residual value are assigned the highest percentile, reflecting the highest predicted constraint (regions with fewer than expected variants). Regions above the 99th percentiles were considered as constrained regions. We used Fisher's exact test to assess if these regions were enriched for tissue- and gene-specific regions and for true positive variants in the curated set of disease-causing non-coding variants from (Caron *et al.*, 2019). We also evaluated enrichment for the controlled genes over Gene Ontology groups, canonical pathways, essential genes, and ClinVar pathogenic genes. For each gene present in GREEN-DB, we selected the highest constraint value across associated regions and evaluated its distribution compared to the gnomAD oe_lof value for the same gene (oe_lof values from https://storage.googleapis.com/gcp-public-data--gnomad/release/2.1.1/constraint/gnomad.v2.1.1.lof_metrics.by_gene.txt.bgz). Finally, we used the Mann–Whitney U test to compare the maximum constraint value between regions controlling genes in the ClinVar pathogenic or essential genes groups and all other regions in GREEN-DB.

*Evaluation of non-coding impact prediction scores*

With the aim of providing a framework useful for variant prioritization, we evaluated the usability of 28 non-coding variant impact prediction scores when applied to WGS data analysis for rare diseases. Among these, we excluded: 10 scores because they do not provide pre-computed values, making them difficult to apply programmatically; 2 scores that were developed specifically for somatic variants; 1 score that provides only disease-specific predictions for a limited set of phenotypes (see Supplementary Table 3). Of the remaining 13 scores, GWAVA and EIGEN provide 3 and 2 different prediction values respectively, for a total of 18 predictors. We compared the performances of these 18 scores when applied to a set of known disease-causing non-coding variants. For this purpose, we used a set of curated disease-associated and neutral variants from (Caron *et al.*, 2019), including 725 true positive examples and 7,250 negative examples. For each score, the evaluation was limited to the subset of scored variants (see Supplementary Table 11). Classification performances were evaluated in R using the ROCR package (Sing *et al.*, 2005), and 3 suggested thresholds for classification were computed: (i) max_ACC: score value achieving maximum accuracy; (ii) TPR90: filtering value corresponding to TPR ≥ 90%; (iii) FDR50: filtering value able to control FDR ≤ 50% with the maximum TPR. For a better representation of the overall classification performances of each score we also computed the overall performance measure (OPM) as described in (Niroula *et al.*, 2015), which is able to summarize all the classification performance metrics when a score is used for filtering purposes.

*Prioritization strategy*

Combining the information in GREEN-DB with population allele frequency, the three best prediction scores (ncER, FATHMM MKL, ReMM), and functional elements (TFBS, DNase peaks, UCNE) we created a prioritization strategy that ranks variants overlapping GREEN-DB regions from Level 1 to Level 4 by summing up evidence of a possible regulatory impact:

1. Level 1: population AF < 0.01
2. Level 2: overlap with at least one functional element among TFBS, DNase peaks or UCNE;
3. Level 3: at least one of the 3 best scores (ncER, FATHMM-MKL, ReMM) with a value above the FDR50 threshold;
4. Level 4: overlaps a GREEN-DB region with constraint value ≥ 0.7.

To further assist in result interpretation, the resulting candidate genes can be ranked based on HPO profiles using GADO (Deelen et al., 2019), and the genes above the 90th percentile in the ranking are suggested as likely disease-related.

*Evaluation on validated disease-causing non-coding variants*

To evaluate the performance of the proposed prioritization method, we applied it to an independent set of curated disease-causing non-coding variants described in (Moyon *et al.*, 2021). We collected the original set of 49 validated variants from https://github.com/DyogenIBENS/FINSURF/tree/master/static/samples and retained for further analysis 45 variants with population AF from gnomAD < 0.01. These variants were selected to be independent from prediction scores training sets in the original paper and are thus also independent from most prediction algorithms compared here. First, we computed the number of variants captured at each of the 4 prioritization levels as defined above.
We then used rare variants from the NA12878 reference sample as a starting point to simulate a WGS patient dataset. The GRCh38 aligned reads for NA12878 sample were downloaded from Illumina Platinum Genomes (https://www.illumina.com/platinumgenomes.html) and deduplicated using Samblaster v0.1.24. Variants were then identified using Deepvariant

v1.0.0 and annotated using vcfanno to include gnomAD / 1000G AF, TFSB / DNase / UCNE overlaps and ncER / FATHMM MKL / ReMM prediction scores. Finally, regulatory annotations from GREEN-DB and prioritization levels were added using GREEN-VARAN. Then, we inserted each validated variant into the WGS variants obtained for the reference sample to generate a simulated disease sample. Using these simulated samples, we first assumed the disease-causing gene was known and evaluated the number of possible candidate variants resulting at each prioritization level as annotated by GREEN-VARAN while restricting to variants linked to the disease genes by our annotations. Then, to better assess the impact of the proposed method in reducing the number of possible candidates, we computed the number of candidate variants resulting from applying it to the whole set of rare WGS variants in each simulated genome, using GADO to prioritize disease-related genes. To be able to run GADO HPO-based prioritization we generated an HPO profile for each simulated genome by selecting a maximum of 5 random HPO terms from those associated with the relevant disease. We then assessed how the known disease gene was ranked by GADO predictions to ensure the approach was effective in prioritizing the correct gene (Supplementary Figure 14). We repeated this analysis assuming either a recessive or dominant mode of inheritance, thus counting only homozygous or heterozygous variants, respectively.

To further test the ability of GREEN-VARAN annotations to capture non-coding variants in distant control elements relevant in human diseases, we evaluated a set of 16 enhancer variants that have been described to have a regulatory effect on disease genes (Supplementary Table 5). For each variant, we evaluated the overlap with GREEN-DB regions and other functional regions (TFBS, DNase, UCNE, dbSuper) and checked whether the affected gene from the original publication is among the ones reported in our database as controlled by the regions overlapping with the variant. Additionally, we assessed if these variants can be classified as “deleterious” based on the FDR50 thresholds we computed for the three best non-coding impact prediction scores. We then assigned to each variant a prioritization level according to the criteria explained above.

*Detection of variants in WGS trios*

To test the impact of our new annotations on the variant prioritization for rare diseases, we applied them to a set of 53 non-consanguineous trios from an internal WGS cohort. For each case, a ranked list of genes potentially relevant for the family phenotype was calculated based on the respective HPO profile using GADO (Deelen *et al.*, 2019) and genes above the 90th percentile in the GADO ranking were selected as best candidates. WGS was performed at a minimum 30X mean coverage, reads aligned to GRCh38 using bwa v0.7.15 (Li and Durbin, 2009), and duplicated reads marked using samblaster v0.1.24 (Faust and Hall, 2014). Small variants were identified from single individual BAM files using deepvariant v1.0.0 (Poplin *et al.*, 2018) and single individual gVCF were merged in a single cohort VCF using GLnexus v1.2.6 with deepvariantWGS optimized settings (Yun *et al.*, 2021). Variants were filtered retaining only variants with quality above 20 and at least 1 individual with GQ $\geq$ 20. The filtered VCF was annotated using SnpEFF v4.3 (Cingolani *et al.*, 2012) and vcfanno

(Pedersen *et al.*, 2016) then our tool was used to integrate non-coding annotations. First, we computed for each individual the number of rare variants (gnomAD / 1000G global population AF < 0.01 and cohort AF < 0.1) and the number of variants overlapping GREEN-DB regions at each of the 4 prioritization levels described in Methods. Then, we computed the number of possible recessive and compound heterozygous candidates when considering in each trio exonic variants only, non-coding variants only, and the combination of both. In detail, for coding variants, we considered 3 distinct groups: all exonic variants, protein-changing variants only, and deleterious variants defined as loss-of-function variants and missense variants with CADD score greater than 20. For non-coding variants, we considered the 4 levels of prioritization defined above and implemented in GREEN-VARAN. For compound heterozygotes we consider combinations involving only potential regulatory variants prioritized at levels 1 and 3, only coding variants in the protein-changing or deleterious groups, and combinations of one regulatory and one protein-changing or coding deleterious variant. In each scenario, we computed the number of candidates considering either all genes, genes prioritized based on the HPO profiles (GADO best candidates as defined above), clinically relevant genes from ClinVar pathogenic list, and PanelApp disease genes.

**Supplementary Results**

*GREEN-DB Database structure*

A schematic representation of GREEN-DB is given in Supplementary Figure 4. The SQLite database contains 16 main tables:

- GRCh37 / GRCh38 regions: GREEN-DB regions coordinate; region type; constraint percentile; closest gene / TSS described by symbol, Ensembl ID and distance; PhyloP100 statistics
- Tissues: tissue(s) where a region or a gene-region interaction is detected
- Genes: controlled gene(s) associated with regulatory regions identified by gene symbol and Ensembl ID. For each interaction, a flag indicates if the region and the connected gene are located in the same TAD
- Methods: method(s) supporting each region and region-gene interaction. This may correspond to the data source when no specific method information was available.
- Phenotypes: potentially associated phenotypes for regulatory regions
- GRCh37 / GRCh38 TFBS: transcription factor binding sites including site coordinates and TF name
- GRCh37 / GRCh38 DNase: DNase hypersensitivity peaks
- GRCh37 / GRCh38 dbSuper: super-enhancers as defined by dbSuper
- GRCh37 / GRCh38 LoF_tolerance: the probability of LoF tolerance for enhancers
- GRCh37 / GRCh38 UCNE: ultraconserved non-coding elements
- GRCh37 / GRCh38 TAD domains: reports TAD domains with a unique ID, genomic coordinates, and the cell line and method of detection.

Main tables (regions, tissues, genes, and methods) are linked by the unique region ID. A unique interaction ID is also assigned to each gene-region pair in the gene table which is linked to methods and tissues tables. Separate tissue tables are included for TFBS and DNase regions describing the cell line/tissue where each region was detected.
Linking tables are included that map the overlap between GREEN-DB region IDs and each of TFBS, DNase, dbSuper, and LoF_tolerance region IDs, reporting also the fraction of overlap. Additionally region_to_TAD and region_to_genes tables are included linking genes and regulatory regions to TAD domains.

*Controlled genes annotation in GREEN-DB*

Overall, ~88% of GREEN-DB regions have a putative association to one or multiple genes, either because these associations were determined experimentally (~32%), or because a gene in close proximity (distance ≤ 10kb, ~56%) can be confidently suggested as a controlled gene (Figure 3A). Considering the 837,879 regions with a validated region-gene association, they interact with a total of 48,230 different genes, covering 67% of all genes and 97% of protein-coding genes from ENCODE v33 basic set. Controlled genes also cover 97, 98, and 100% of clinically relevant genes from PanelApp (Martin *et al.*, 2019), ClinVar (pathogenic genes only), and the ACMG actionable genes list, respectively (Supplementary Table 6).
Distal control elements (silencers, enhancers, bivalents) are mostly located outside the controlled gene, balanced between up- or downstream locations, while most of the promoter regions are located inside genes or upstream of them (Figure 2B). As expected, the distance between a region and its controlled gene(s) is larger for enhancers and silencers, which appear to be mostly located from about 10kb up to several Mb away from their controlled gene (Figure 2C). When we analyzed the relationship between GREEN-DB regions and their controlled genes (taking only experimentally-associated genes into account), we saw that the closest gene is among annotated controlled genes only for ~70% of enhancers and ~25% of silencers, while this proportion is much higher (~90%) for promoters, as expected. Even when the closest gene is controlled, it is the only associated gene in just 24% and 5% of cases for enhancers and silencers, respectively. Interestingly, even when considering only GREEN-DB regions located within a gene, this gene is among the controlled ones in less than 50% of cases for enhancers, silencer, and bivalent regions (Figure 3D).
The region-to-gene relationship showed a high degree of specificity, with most regions controlling less than 5 genes, while several genes are controlled by multiple regions (Supplementary Figure 6). Regions active in multiple tissues usually control more genes, suggesting a tissue-specific region-to-gene relationship (Supplementary Figure 7).
Finally, gene-set enrichment analysis performed on the 491 genes with an extremely large regulatory space showed that these genes are strongly enriched for essential genes derived from mouse studies (p-value 7.88E-97, FDR 1.11E-91) as well as genes involved in developmental processes, cell differentiation, and other essential biological functions (Supplementary Table 16).

*Pre-processed datasets available in the GREEN-DB collection*

To facilitate non-coding annotations in standard bioinformatics pipelines we also provide, in addition to GREEN-DB regions tables, a set of standard tables in BED-like format representing the 6 additional datasets in Supplementary Table 2. These tables have genomic coordinates in the first 3 columns, followed by additional information. We also prepared similar pre-processed tables for the 18 prediction scores listed in Supplementary Table 12, adding reference and alternate alleles in columns 4 and 5 when available, and then the score value. These tables are provided compressed and indexed and freely accessible from Zenodo repositories (see https://green-varan.readthedocs.io/en/latest/Download.html)

*The GREEN-VARAN tool and annotations*

To facilitate variant annotation using GREEN-DB, we have developed a companion tool (GREEN-VARAN) that processes any VCF containing small variants and adds regulatory annotations from the GREEN-DB (see the table below). The tool is freely available from the GitHub repository https://github.com/edg1983/GREEN-VARAN.

Besides regulatory region annotations, the tool can also add a prioritization level for each variant, integrating GREEN-DB annotations with population AFs, overlap with additional relevant regions, and prediction scores values. The provided configuration JSON file reproduces the 4 level prioritization strategy described in Methods. However, the configuration can be modified to also include additional functional regions or annotated float values allowing the user to tweak the prioritization exploiting additional annotations, such as functional regions determined by specific ChIP-seq or ATAC-seq experiments. If the user provides a list of genes of interest, GREEN-VARAN can add a tag into VCF INFO marking regulatory variants potentially relevant for those genes based on GREEN-DB annotations. Finally, the tool can update the gene consequence description field generated by popular software like SnpEFF or bcftools CSQ, adding new gene annotations based on the GREEN-DB regulatory annotations and setting a variant consequence according to the regulatory region type. This simplifies integration with downstream segregation analysis tools or gene burden tests that usually rely on gene consequence annotation fields.

The following table summarizes the annotation fields added by GREEN-VARAN:

| Annotation tag | Data type | Description |
|---|---|---|
| greendb_id | String | Comma-separated list of GREEN-DB IDs identifying the regions that overlap this variant |
| greendb_stdtype | String | Comma-separated list of standard region types as annotated in GREEN-DB for regions overlapping the variant |
| greendb_dbsource | String | Comma-separated list of data sources as annotated in GREEN-DB for regions overlapping the variant |

| | | |
|---|---|---|
| greendb_level | Integer | Variant prioritization level computed by GREEN-VARAN based on the supporting evidences as configured in the configuration file |
| greendb_constraint | Float | The maximum constraint value across GREEN-DB regions overlapping the variant |
| greendb_genes | String | Possibly controlled genes for regulatory regions overlapping this variant |
| greendb_VOI | Flag | If a list of genes of interest is provided, this flag is set when any of the input genes is among the possibly controlled genes for overlapping regulatory regions. |

The tool can eventually accept an additional list of values or region flags to be evaluated from the INFO field when computing the prioritization level, thus allowing easy integration of additional dataset-specific annotations, like TF binding sites derived from specific ChIP seq experiments. In this case the prioritization level is increased for each of the additional regions or values.
The GREEN-VARAN tool can also be used to annotate structural variant VCF adding the same annotations described for small variants, except the prioritization level which can not be computed in this case. Finally, the tool can be used to process a previously annotated VCF and extract a table of variant IDs and corresponding GREEN-DB region IDs that can be used as input for GREEN-DB query to automatically retrieve additional details on the regulatory regions from the database.
We also provide a query interface for the database that, given a list of variants, region IDs or a table produced by GREEN-VARAN, can be used to extract detailed information on the relevant regulatory regions, controlled genes and overlapping functional regions including the tissue of activity.

*Nextflow annotation workflow*
To be able to annotate prioritization levels as described in the Methods section, GREEN-VARAN needs a set of additional information to be already annotated in the input VCF file. These include population allele frequencies (such as AF from gnomAD or 1000G); overlap with functional regions, namely transcription factor binding sites, DNase peaks and ultraconserved elements; and the 3 best prediction scores, namely ncER, FATHMM MKL and ReMM. The relevant INFO tags for these annotations can be configured as needed in the configuration file. To facilitate end-to-end prioritization of non-coding variants, we also provide a Nextflow workflow based on vcfanno annotation tool (Pedersen *et al.*, 2016). Given a VCF containing small variants from WGS, this workflow can be used to automatically add all the annotations needed by GREEN-VARAN and then perform regulatory variants annotation and prioritization. If not available locally, the datasets needed

for annotations will be automatically downloaded from the corresponding repository of pre-processed data (see pre-processed dataset section).

### Supplementary Figures

**Supplementary Figure 1.** ***Random regions used as control regions***

We generated a set of control regions by randomly sampling across the genome the same number of regions present in GREEN-DB. Random regions have a similar distribution across chromosome (A) and a similar size distribution (B, zoom in C).

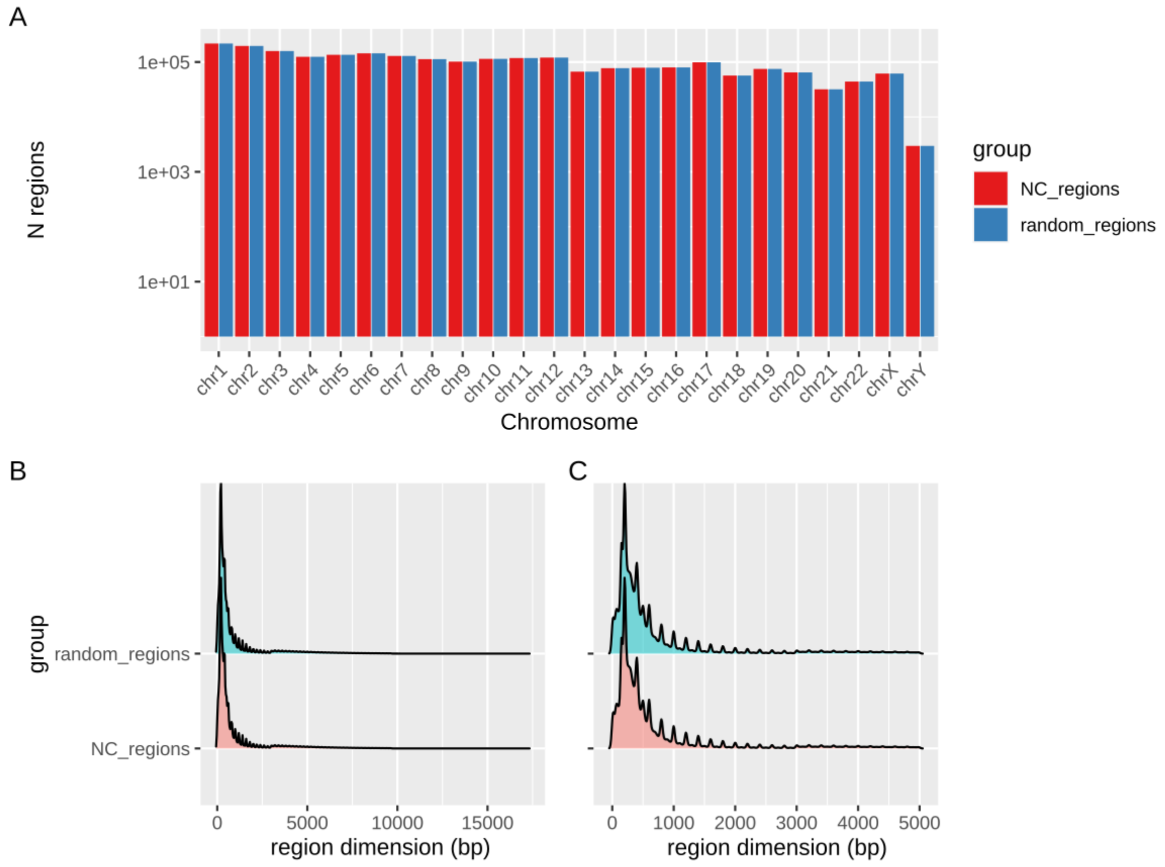

**Supplementary Figure 2.** ***Distribution of regions across chromosomes***
Count of regions in GREEN-DB across human chromosomes, stratified based on region type.

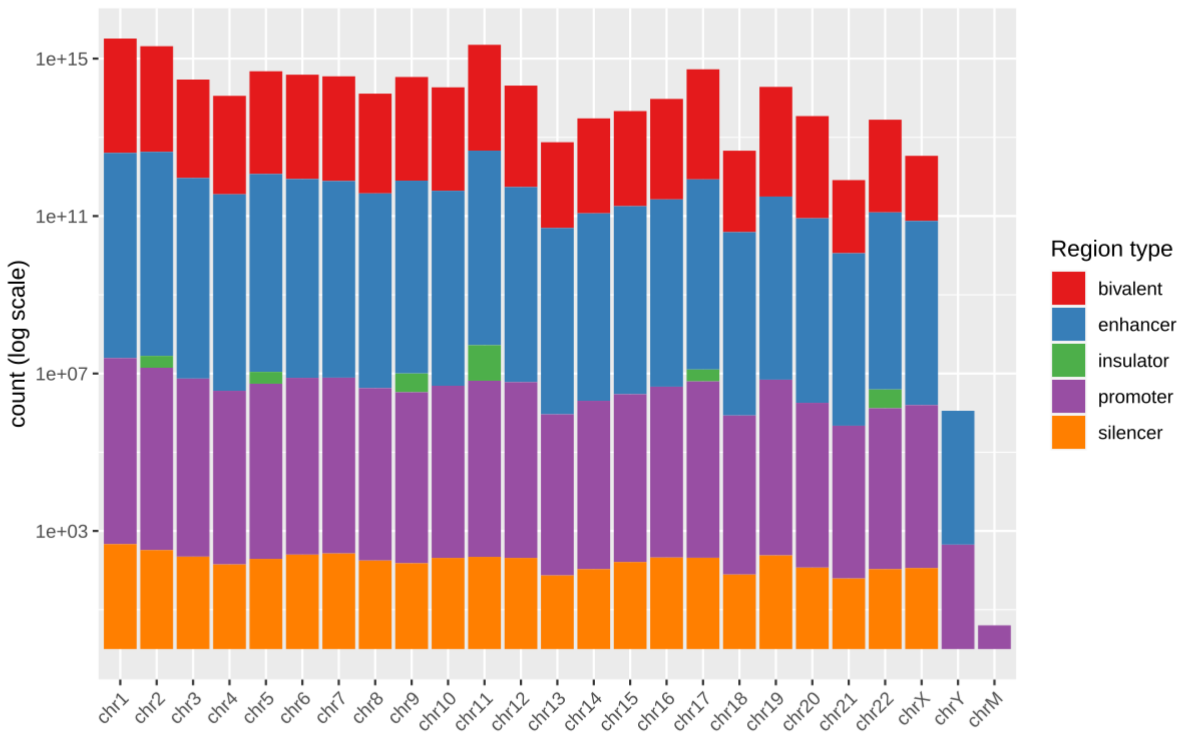

**Supplementary Figure 3.** ***Contribution of different genomic regions to GREEN-DB regions***

We calculated the fraction of bases overlapped by different genomic regions (intergenic, intronic, exonic, CDS, UTR) for all regions in GREEN-DB as well as for each region type separately. Gene definitions are based on the GENCODE v33 basic set.

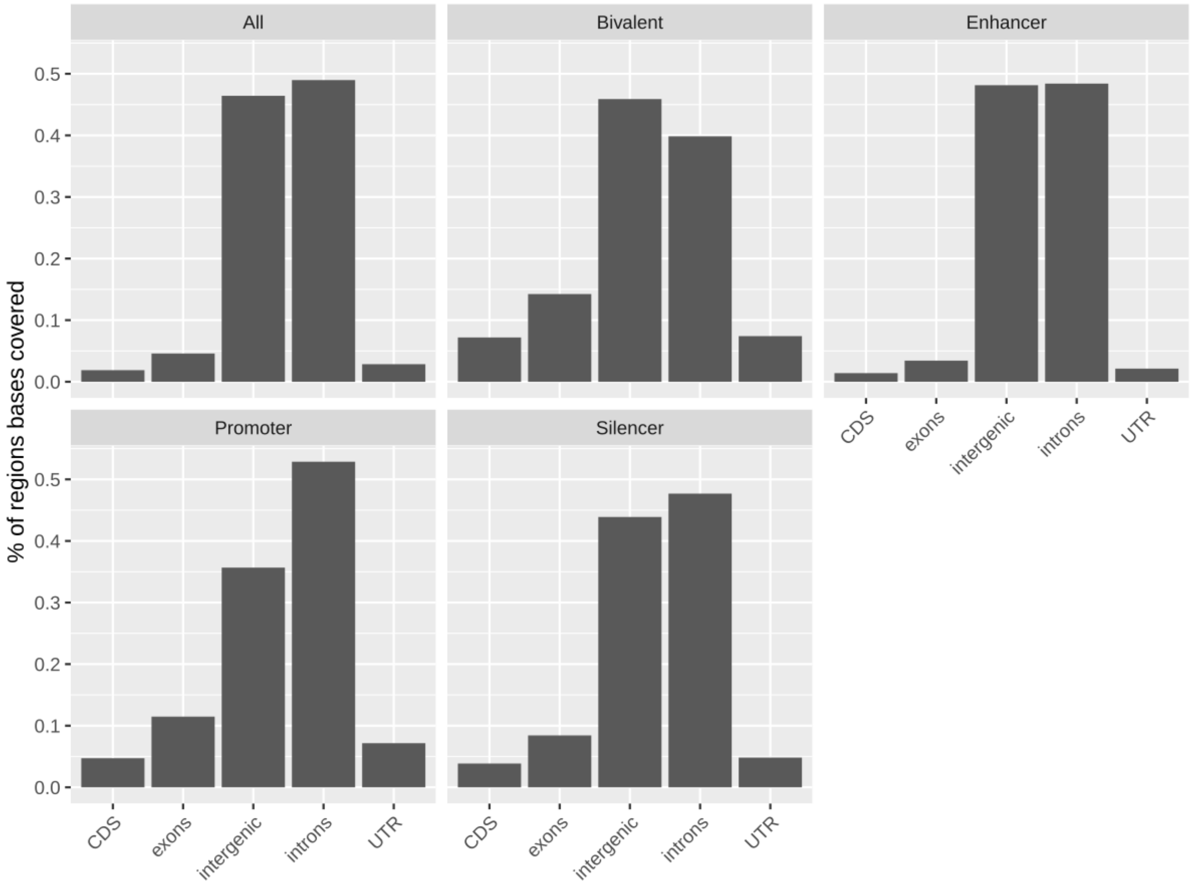

**Supplementary Figure 4.** ***Schematic representation of GREEN-DB***

The main structure of the GREEN-DB SQlite database is represented to illustrate the information contained in each table and how the tables are connected together. Genomic coordinates for both GRCh37 and GRCh38 are provided in distinct tables for the regulatory regions and the additional datasets.

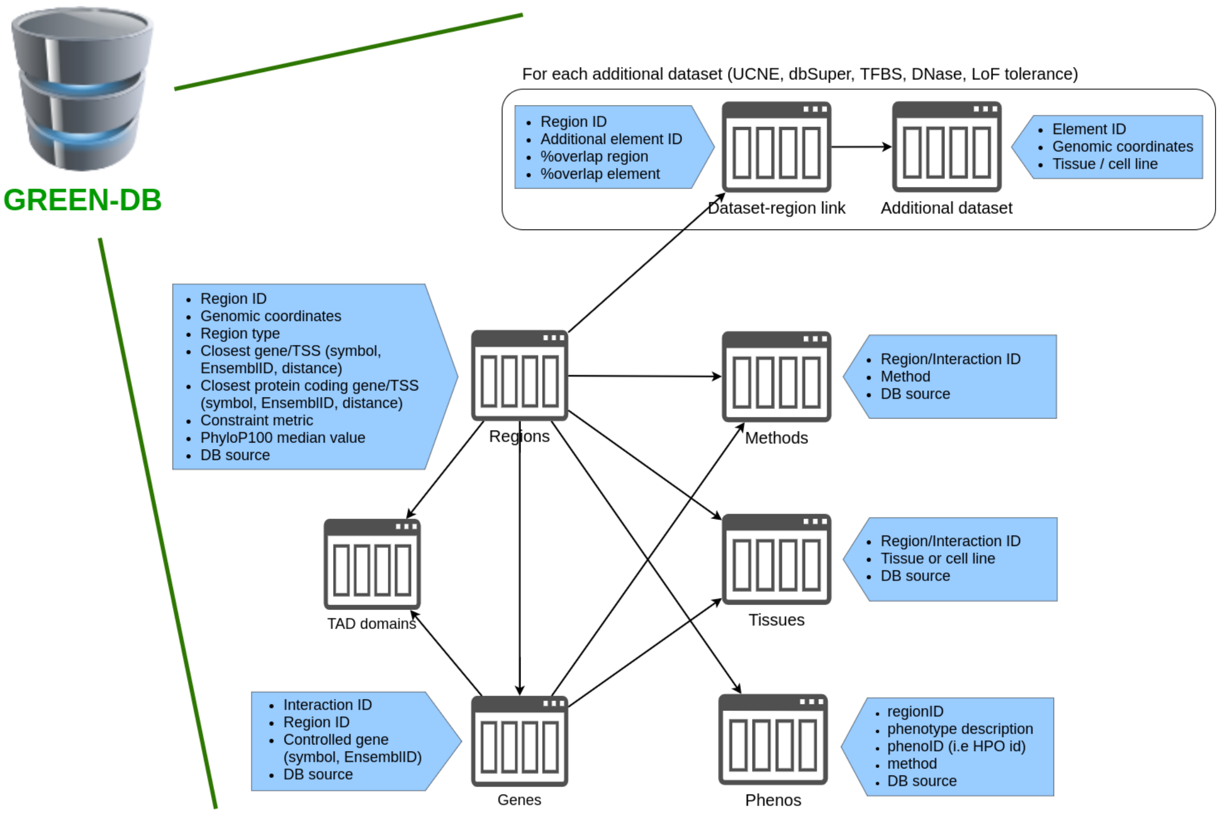

**Supplementary Figure 5.** ***Evaluation of the GREEN-DB regions***
(A) Using Fisher's exact test, we assessed the presence of an enriched overlap between regions collected in the GREEN-DB and: DNase HS peaks (Dnase), transcription factor binding sites (TFBS) from ENCODE, ultraconserved non-coding elements (UCNE), significant eQTLs from GTeX v8 (GteX eQTLs), a curated set of disease-causing non-coding variants (TrueSet Vars), segmental duplications (SegDup) and low-complexity regions (LCR). (B) GREEN-DB regions have a higher proportion of highly conserved bases per region at either 1, 1.5, or 2 PhyloP100 thresholds compared to random regions. (C) GREEN-DB regions also showed higher per region median and maximum score for FATHMM MKL, ncER and ReMM.

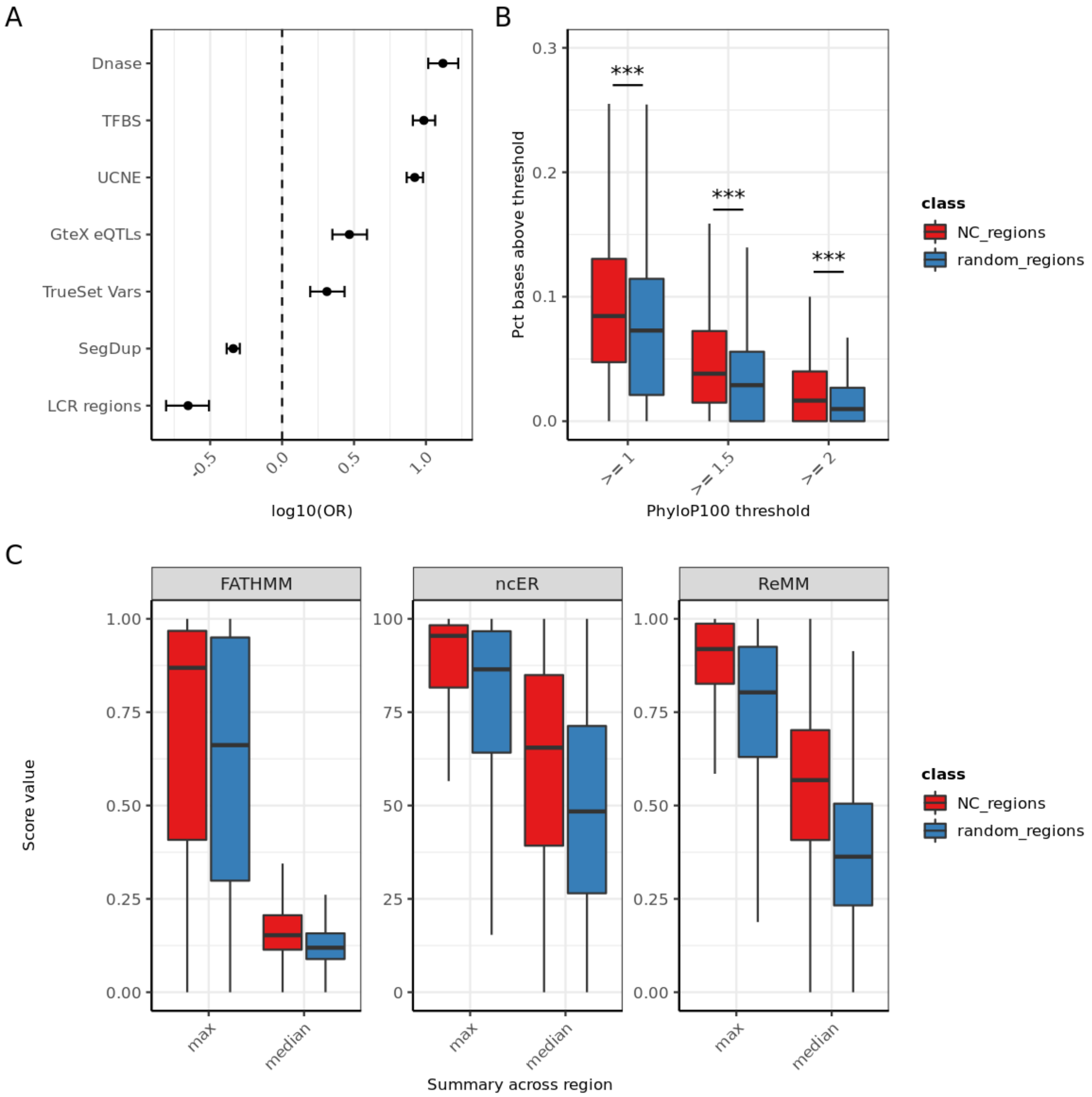

**Supplementary Figure 6.** ***Summary of gene-region connections***
Based on the 839,807 regions with a validated gene association in GREEN-DB, we calculated the number of genes controlled by each region (A) and the number of regions controlling each gene (B).

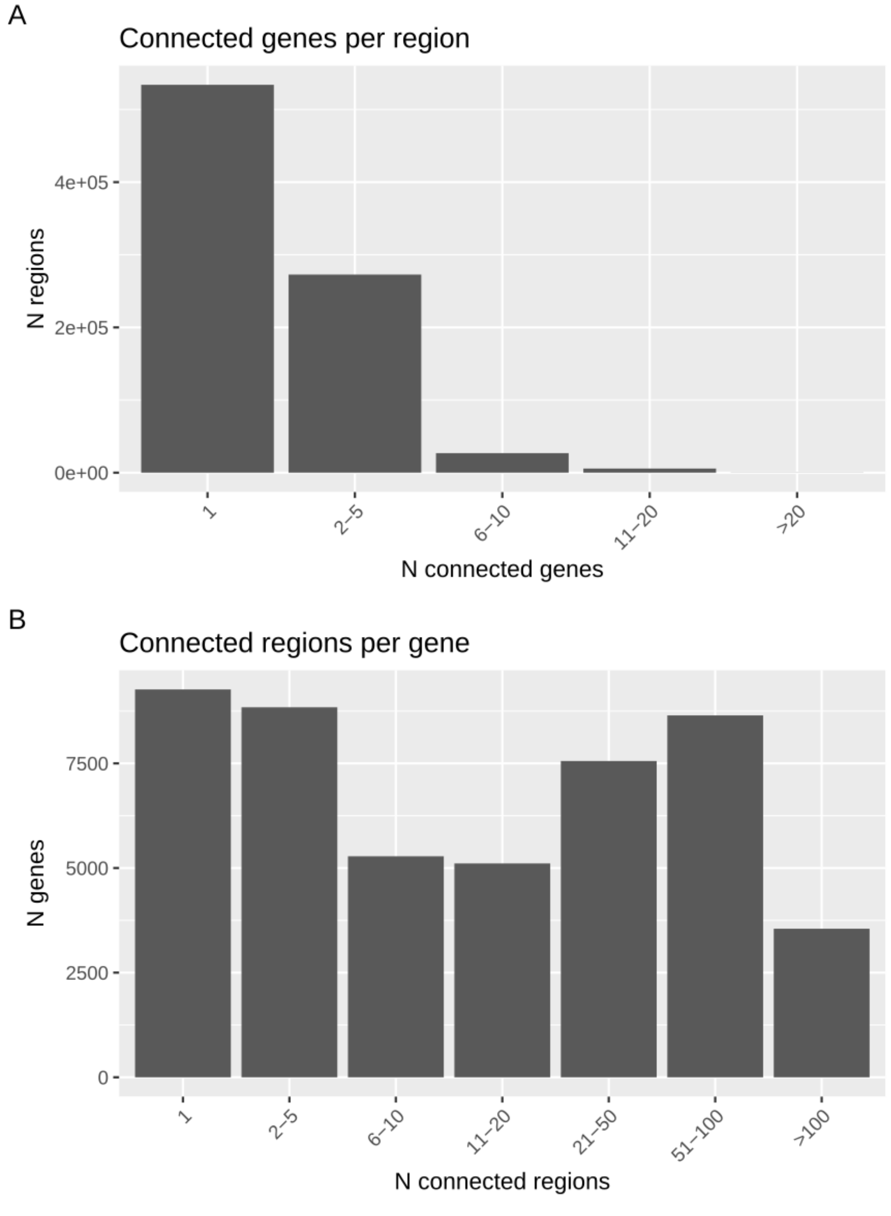

**Supplementary Figure 7.** ***Tissue of activity and its relationship with N controlled genes***
Based on the information from GREEN-DB we calculated the number of tissues in which each region is potentially active (A), showing a large proportion of tissue-specificity. Considering the 839,807 regions with a validated gene association in GREEN-DB, we then evaluated the relationship between the number of tissues and the number of controlled genes for each region (B). Number of tissues correlates with number of controlled genes (blue line). To avoid biases due to the fact that tissue information may be poorly annotated for some regions, we repeated the same analysis considering the median N tissues (red diamonds) across regions controlling a certain number of genes. This resulted in a stronger correlation (red line).

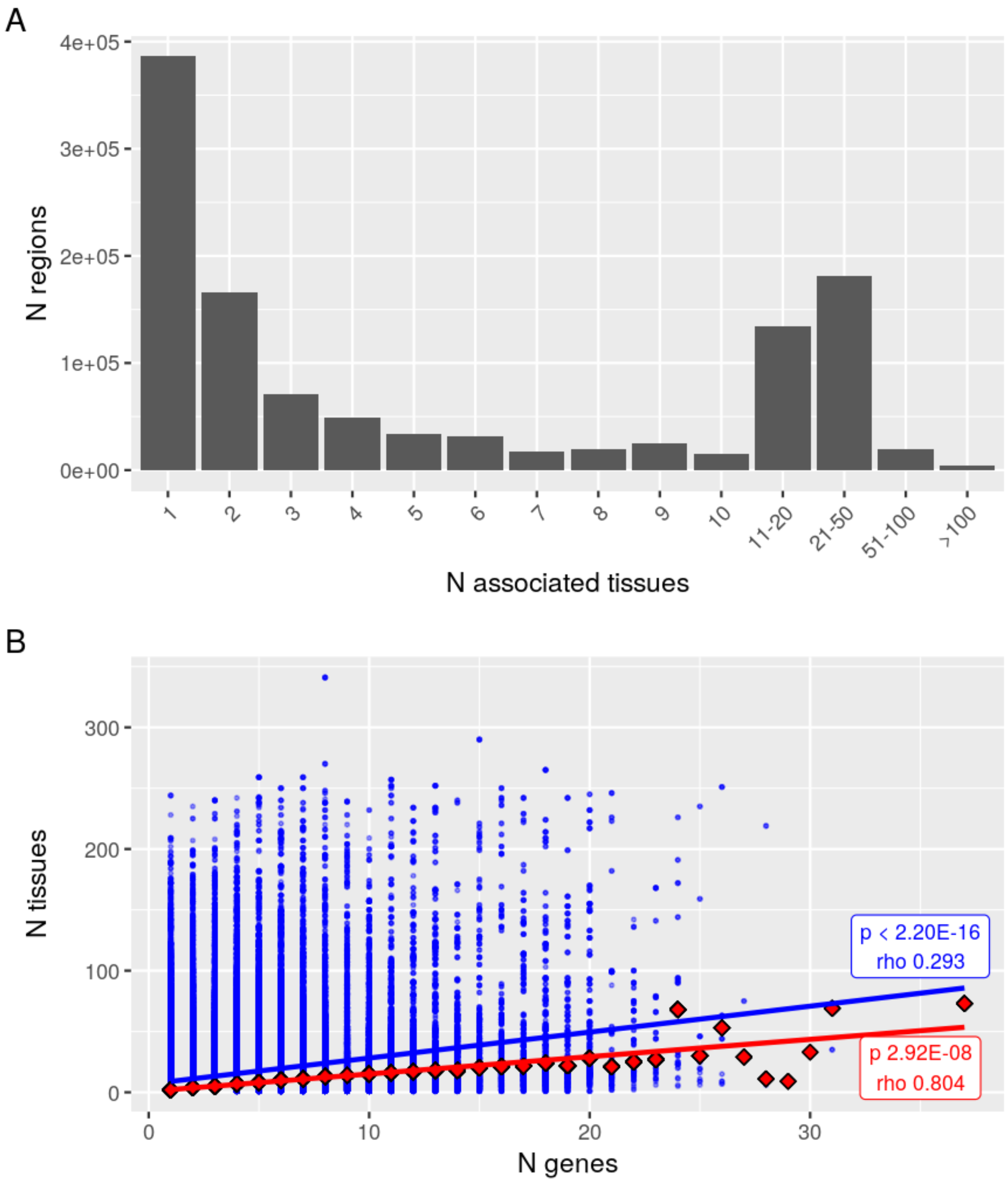

**Supplementary Figure 8.** ***Identification of constrained GREEN-DB regions***
(A) Using data from gnomAD v3, we calculated the number of variants across each region and constructed a model of N variants dependent on sequence length, %GC, overlap with segmental duplication and LCR regions. (B) We then used the residuals from this model to rank GREEN-DB regions, selecting as constrained those in the 99th percentile of low residual values (dashed line). (C) These constrained regions are more conserved than other regions in GREEN-DB, showing a higher fraction of bases with PhyloP100 values >= 2, and they are mostly represented by enhancer and promoters (D).

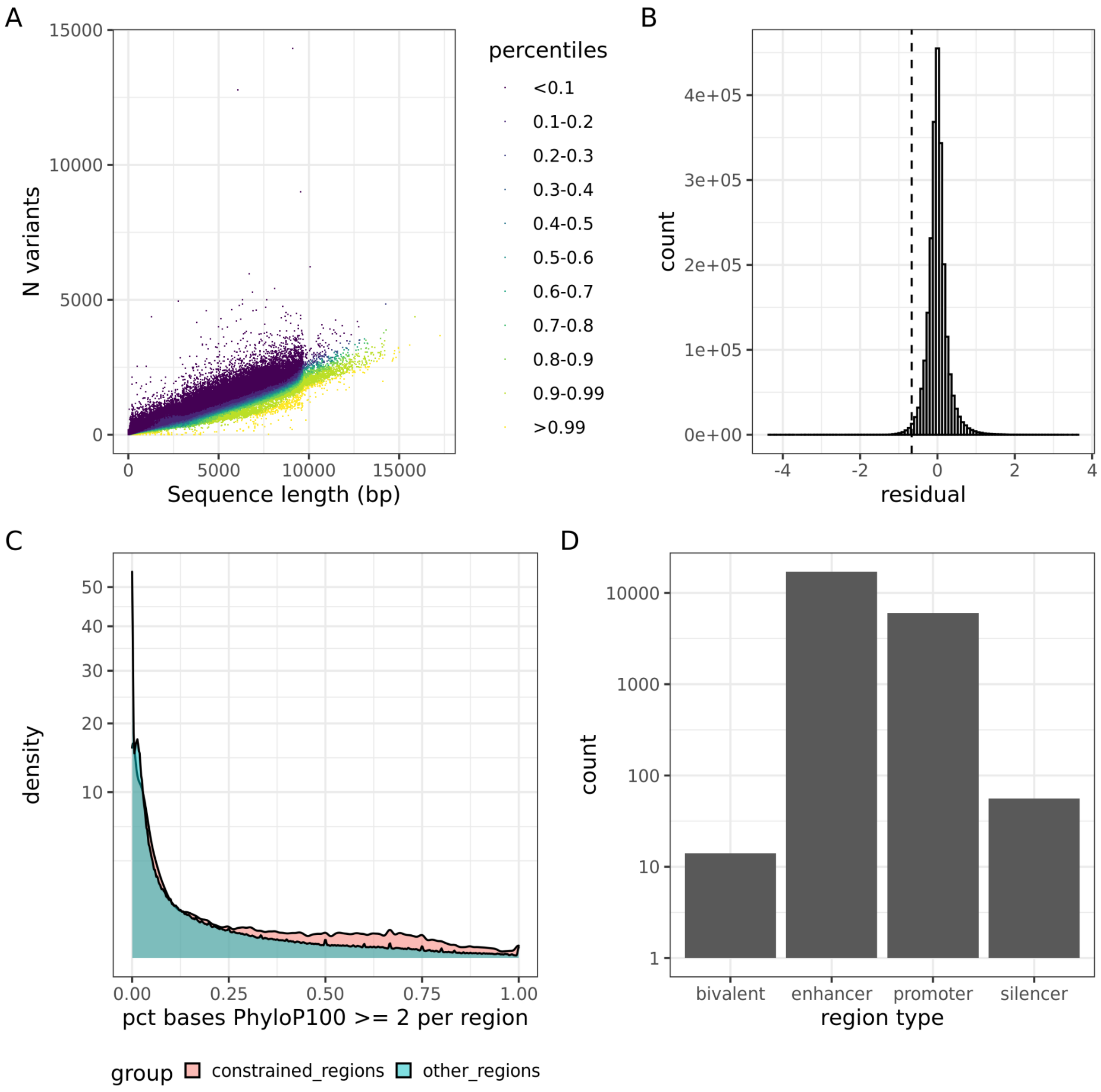

**Supplementary Figure 9.** ***Tissue and gene-specificity of GREEN-DB constrained regions***
We compared the tissue (A) and gene (B) specificity of GREEN-DB constrained and not constrained regions, counting the regions active in a single tissue or associated to a single gene. The reported p value was computed by Fisher's exact test for enrichment of tissue-/gene-specific regions among  GREEN-DB constrained regions.

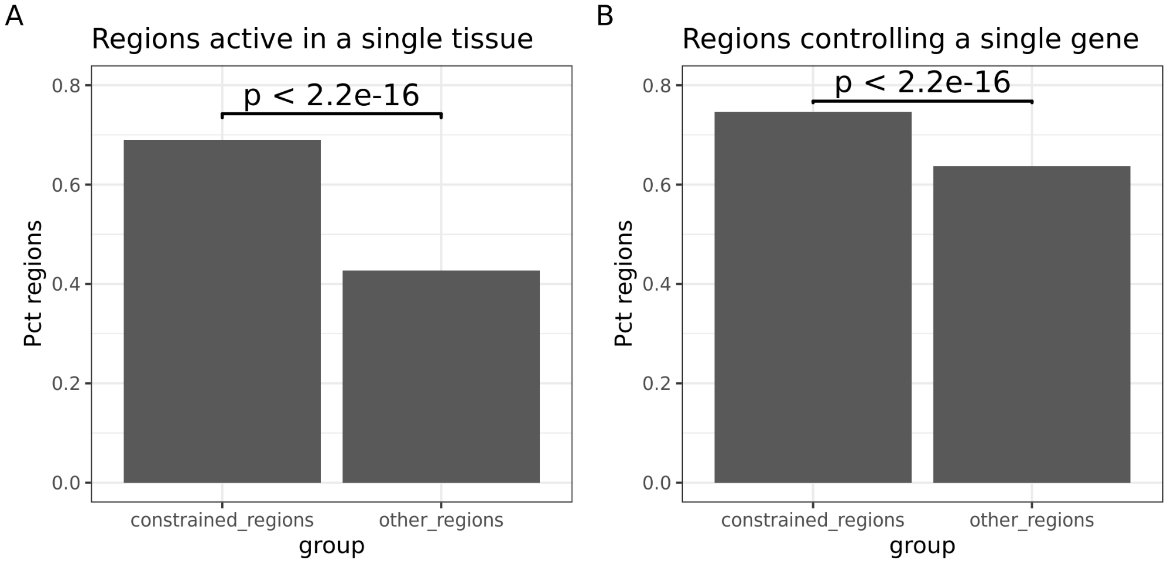

**Supplementary Figure 10.** ***Relationship between constrained GREEN-DB regions and essential genes.***

For each gene reported in GREEN-DB, we considered the maximum constraint value across the associated regions and compared the distribution of these values between general genes and genes in the essential genes (A) or ClinVar pathogenic (B) groups. Both groups appear to be controlled by regions with higher constraint. However, the median constraint value considering all associated regions is only slightly higher for essential / pathogenic genes compared to normal genes (C and D)

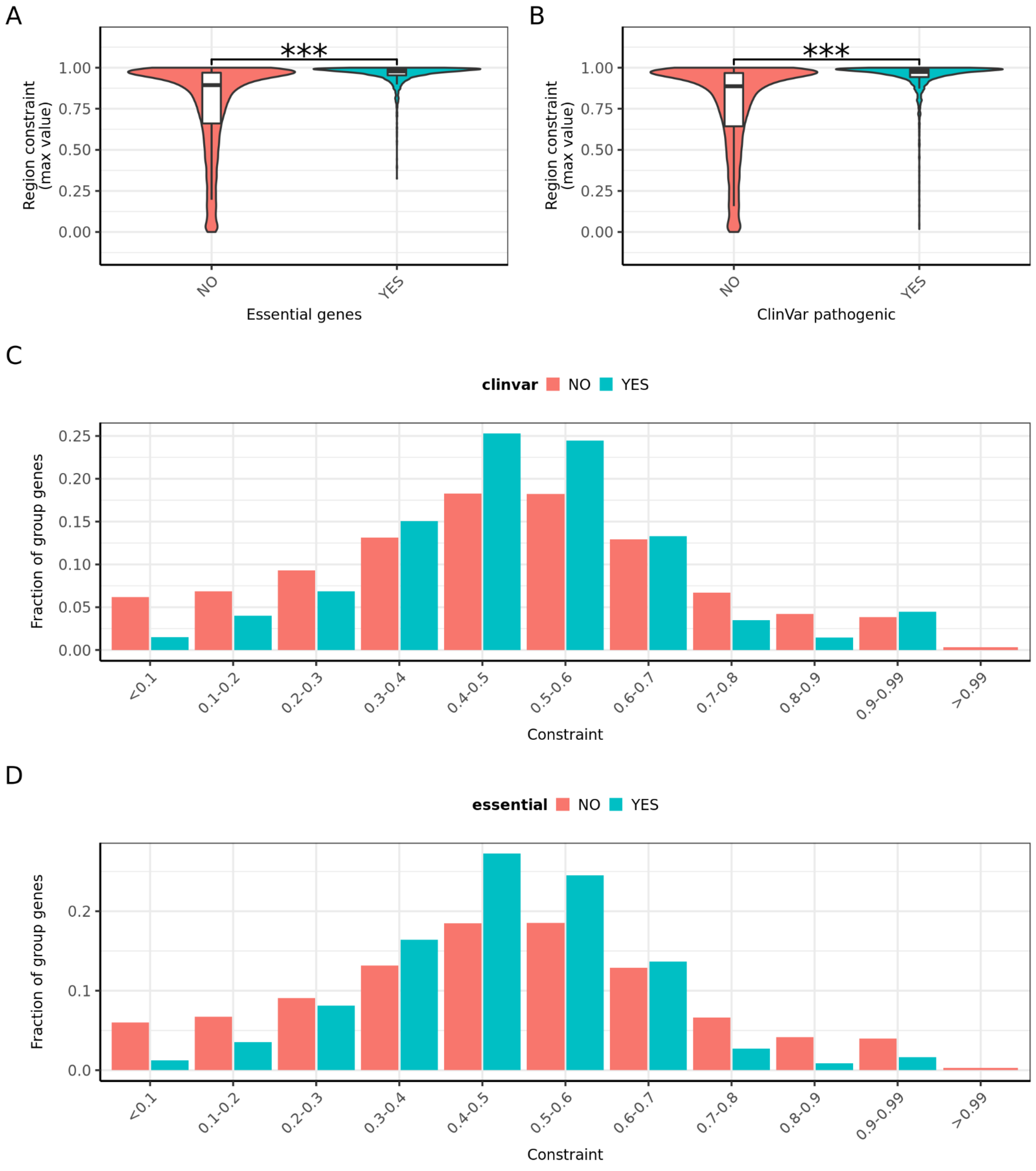

**Supplementary Figure 11.** ***Performances of the non-coding prediction scores***
We tested 18 different non-coding prediction scores using a curated set of non-coding disease-associated variants. Performances were evaluated using (A) ROC curves and (B) precision-recall curves. AUC values are reported for each score in (A).

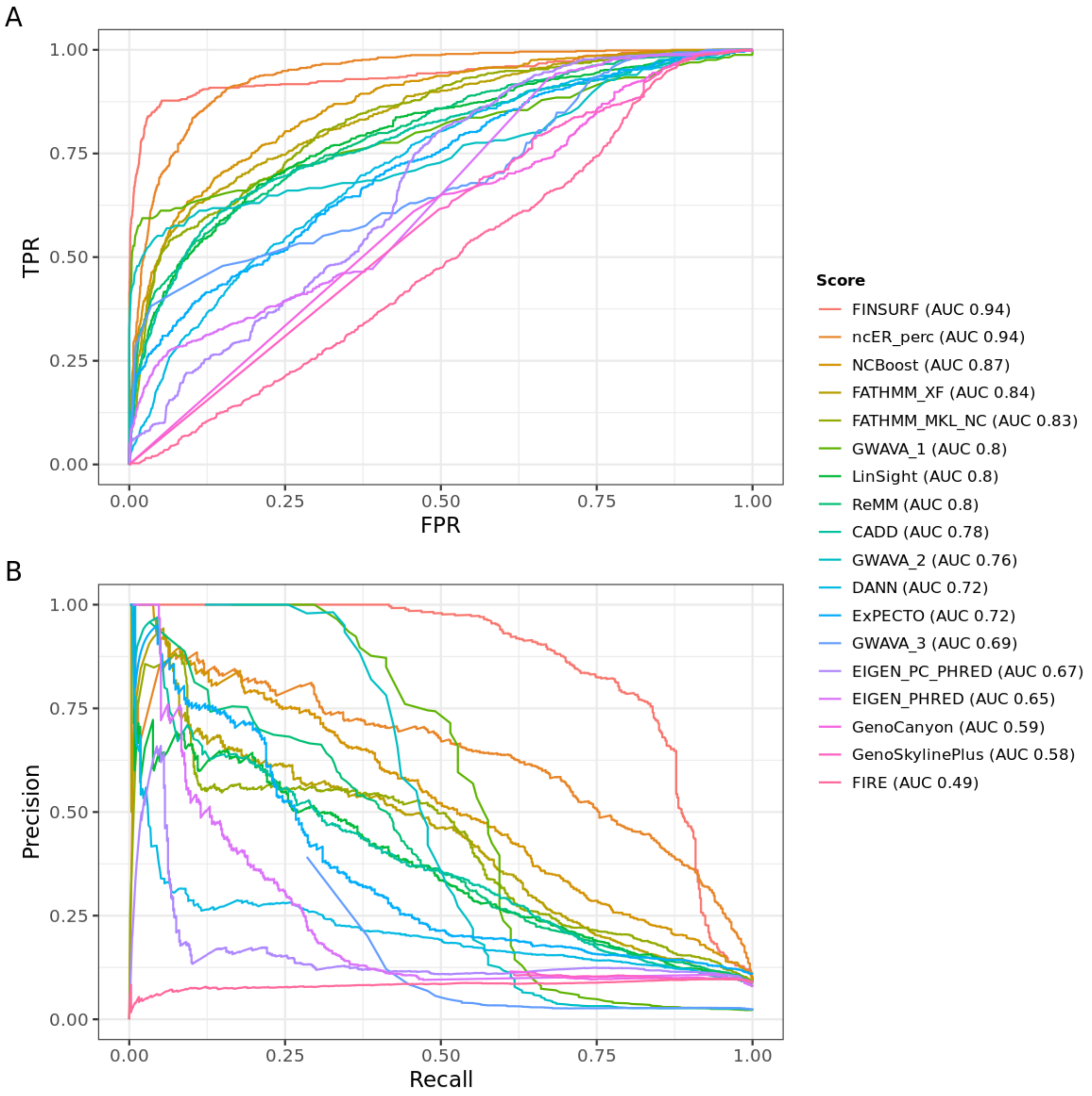

**Supplementary Figure 12.** ***TPR-TNR curves and suggested classification thresholds***
For each score, we computed TPR-TNR curves. In each curve we represented 3 points corresponding to the FDR50, TPR90 and MAX_ACC thresholds calculated as described in the Methods section.

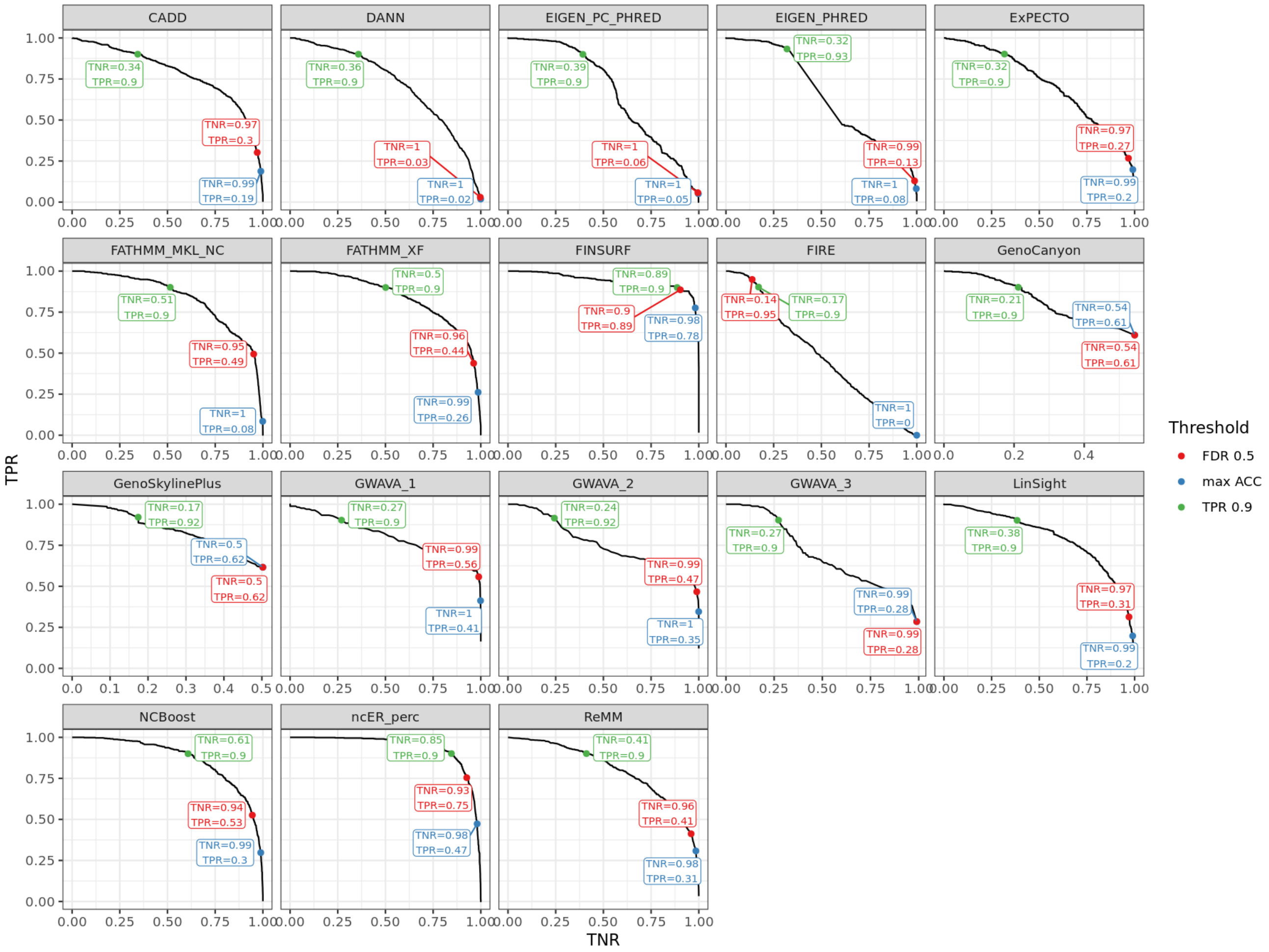

**Supplementary Figure 13.** ***Number of variants overlapping GREEN-DB from WGS trio analysis***

Distribution of the number of variants overlapping GREEN-DB regions detected across 53 WGS trios. Counts are reported for all rare variants, as well as for rare variants overlapping GREEN-DB regions and prioritized in one of the 4 levels described in methods.

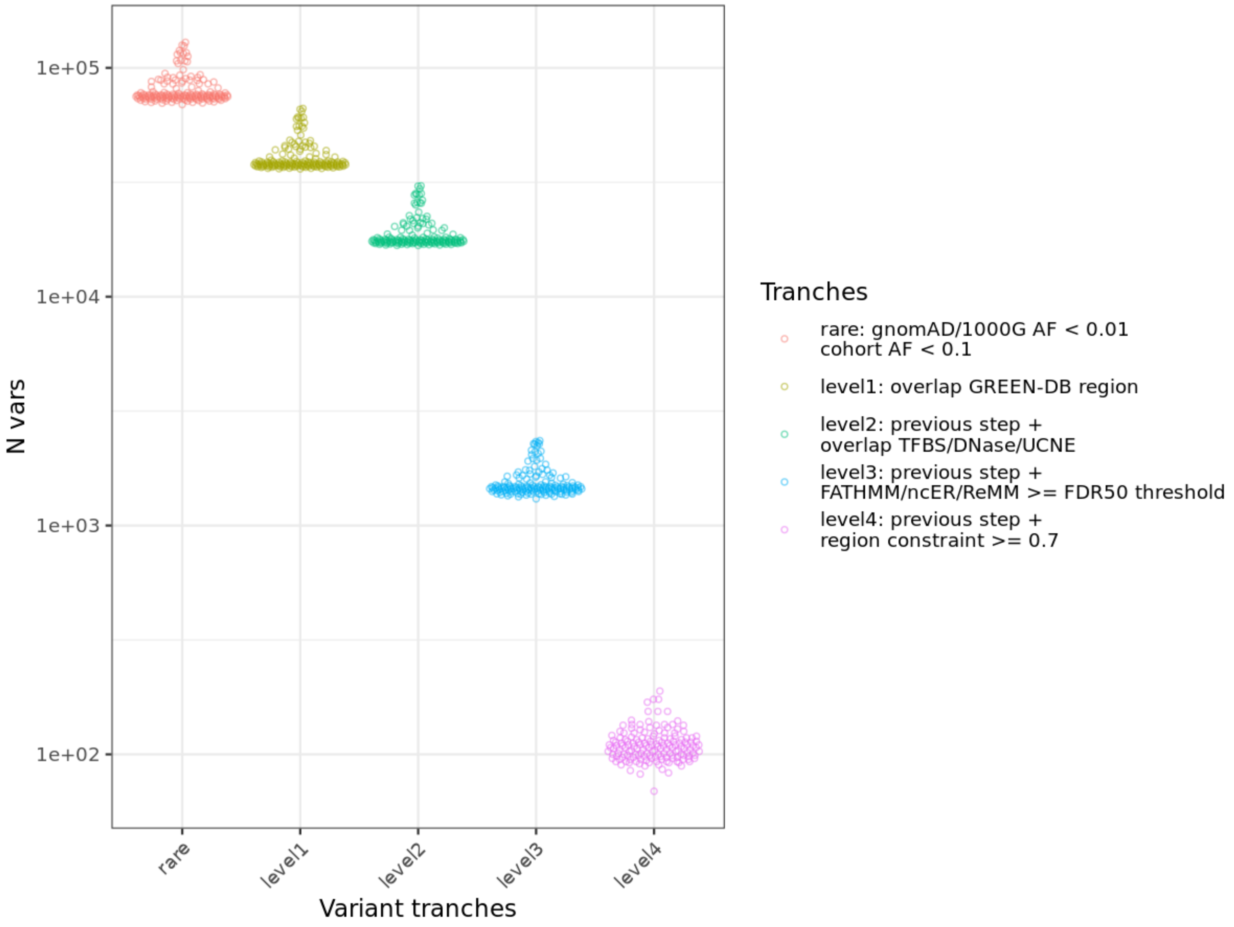

**Supplementary Figure 14.** ***HPO based prioritization for profiles based on disease genes corresponding to the 45 validated variants evaluated in the study***

For each of the 45 disease-causing validated variants from (Moyon *et al.*, 2021), we extracted from the HPO ontology all the HPO terms associated with the corresponding disease (N_hpo). We then generated for each disease a simulated profile subsampling these HPOs to a maximum of 5 terms and evaluated how the causative gene was ranked by GADO using the corresponding profile (GADO_rank). The corresponding percentile in the GADO ranking is shown in the bottom panel (GADO_percentile)

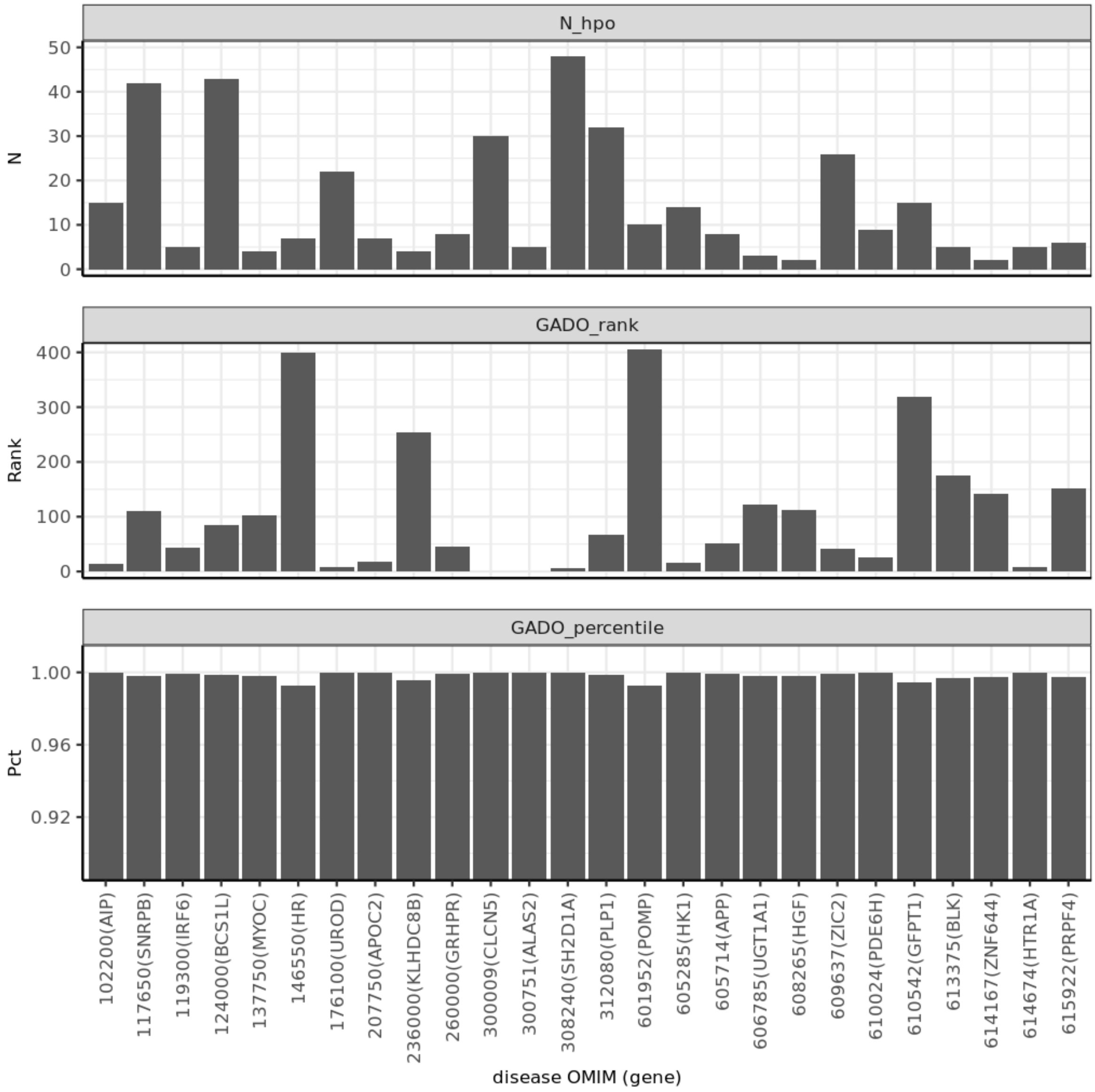
